# Systematic Analysis of Autophagy Identifies Atg9 Vesicles as the Origin of the Phagophore

**DOI:** 10.1101/2022.08.03.502680

**Authors:** David Broadbent, Carlo Barnaba, Jens C. Schmidt

## Abstract

Autophagy is a catabolic pathway required for the clearance and recycling of cytoplasmic materials. Upregulation and dysfunction of autophagy contributes to the pathology of cancer and neurogenerative diseases, respectively. To define the molecular mechanisms that control autophagic flux it is critical to quantitatively characterize the dynamic behavior of autophagy factors in living cells. Using a panel of 9 cell lines expressing HaloTagged autophagy factors from their endogenous loci, we systematically analyze the abundance, single-molecule dynamics, and autophagosome association kinetics of a wide variety of autophagy proteins involved in the initiation and maturation of the autophagosome. Our results reveal that phagophores are initiated by the accumulation of autophagy factors on mobile ATG9 vesicles and tethering of these ATG9 vesicles to donor membranes by ATG2 is a key step in phagophore maturation. In addition, we demonstrate that the overall lifetime of an autophagosome is approximately 160 seconds and the majority of phagophore initiation events fail to produce mature autophagosomes. In total our work establishes a new experimental framework to quantitatively analyze autophagy and demonstrates that ATG9 vesicles are the seeds for autophagosome formation in human cells.

## Introduction

Autophagy is a conserved catabolic process that recycles damaged organelles and protein aggregates, or nonspecifically degrades cellular material to provide nutrients for cell proliferation, particularly when cells face chemical stress or nutrient starvation (White, 2015; Yu et al., 2018). The hallmark of autophagy is the formation of double-membrane autophagosomes, which sequester cargo and then fuse with lysosomes to trigger the degradation of their contents. Alterations of autophagy have been implicated in the pathology of several human diseases. During the aging of human cells, key autophagy factors are reduced in abundance resulting in the downregulation of autophagic flux which increases the susceptibility to the two most common neurodegenerative disorders, Alzheimer’s and Parkinson’s disease (Lu et al., 2020; Nixon, 2007). In contrast, upregulation of autophagy has been shown to promote a variety of cancers by providing nutrients for rapid tumor proliferation (Gewirtz, 2014).

The life cycle of autophagosomes encompasses four distinct steps: initiation, maturation, closure, and degradation. Autophagosome formation can be initiated nonspecifically (canonical autophagy) or by a target, for instance, a damaged organelle, that requires degradation (Kirkin, 2020; Lamb et al., 2013). Canonical autophagy is induced under starvation conditions or by chemical stress and begins with the formation of the phagophore, a rapidly expanding double-membrane that will eventually become an autophagosome (Lamb et al., 2013). Importantly, the molecular nature of the phagophore is hotly debated. Initially, it was suggested that specific regions of the endoplasmic reticulum (ER) are remodeled into the phagophore (Ge et al., 2017). More recently, a growing body of evidence indicates that the phagophore is formed by ATG9-containing vesicles, which expand to form a mature autophagosome (Chang et al., 2021a; Chang et al., 2021b). ATG9 is the only known autophagy-related protein that contains a transmembrane domain and has recently been shown to have lipid scramblase activity, which facilitates the exchange of phospholipids between the outer to the inner leaflets of the membrane it is embedded in (Matoba et al., 2020; Matoba and Noda, 2020; Noda, 2021). For this reason, ATG9 vesicles are prime candidates for the origin of the phagophore; as ATG2 transfers lipid to the outer leaflets, ATG9 equilibrates the membrane, growing a vesicle into an autophagosome (Matoba et al., 2020; Matoba and Noda, 2020; Noda, 2021). Regardless of the membrane structure autophagosomes originate from, canonical autophagy is initiated through a complex phosphorylation cascade by the Unc-51 like autophagy activating kinase (ULK1/2) complex 1. Besides Ulk1/2 isoforms, the Ulk1-kinase complex comprises the focal adhesion protein FIP200, a HORMA domain-containing protein ATG13 that bridges the interaction between ULK1, FIP200, and ATG101 (Ganley et al., 2009; Mercer et al., 2009; Shi et al., 2020). The ULK1 substrates include many downstream autophagy factors, and these phosphorylation sites serve as key switches for autophagy protein recruitment to the phagophore (Ganley et al., 2009; Mercer et al., 2018; Mizushima, 2010). One essential step that drives autophagosome maturation is the activation of the PI3K complex at the phagophore, which modifies phospho-inositol lipids leading to a local enrichment of PI3P (Mizushima, 2010). The presence of PI3P is sensed by the WIPI3/4 proteins, which subsequently recruit the lipid transferase ATG2 to the growing autophagosome (Chowdhury et al., 2018; Dooley et al., 2014; Otomo et al., 2018). In addition to ATG2, WIPI1-4 also recruits the ATG5-ATG12-ATG16 complex (Fracchiolla et al., 2020; Lystad et al., 2019), which acts similarly to ubiquitin ligase complexes but instead conjugates LC3/GABARAP proteins to phosphatidylethanolamine at the autophagosome membrane (Kirisako et al., 1999; Shpilka et al., 2011). Once conjugated to the autophagosomal membrane, LC3 and GABARAP proteins serve as anchors to tether cargo targeted for degradation to the autophagosome (Schaaf et al., 2016). Finally, when maturation and cargo sequestration are complete, the autophagosome closes into double-membrane vesicles and fuses with the lysosome to trigger the degradation of its contents (Berg et al., 1998; Nakamura and Yoshimori, 2017). While we have detailed knowledge of the biochemical activities of many autophagy factors, how their dynamic recruitment to the phagophore regulates autophagosome formation and thereby overall autophagic flux has not been analyzed in great detail. To identify the ideal intervention points to inhibit or upregulate autophagy as a therapeutic strategy to treat cancer and neurological disorders, respectively, it is critical to quantitatively define the molecular mechanisms underlying autophagosome formation.

In this study, we establish a collection of cell lines using genome editing that express HaloTagged autophagy proteins involved in initiation and maturation of autophagosome formation, from their endogenous locus. This approach maintains the expression levels of the tagged autophagy factors and retains all regulatory mechanisms conferred by the endogenous genomic locus of the respective gene. The HaloTag is a versatile protein tag that can be covalently linked to cell-permeable ligands, which facilitates fluorescent labeling, targeted protein degradation, and pulse-chase experiments. Using our collection of cell lines, we systematically quantify the absolute abundance, single-molecule diffusion dynamics, and the autophagosome recruitment kinetics of these autophagy proteins. Our live cell single-molecule imaging experiments reveal that the initiation of autophagosome formation is locally controlled, rather than by cell-wide changes. Strikingly, our results demonstrate that ATG9 is the only autophagy factor that is not locally enriched at sites of autophagosome formation, consistent with ATG9 vesicles containing a limited number of ATG9 molecules forming the seeds for phagophore formation. Our systematic analysis of autophagy factor foci kinetics revealed that the average lifetime of an autophagosome is approximately 160 seconds. Within these autophagosomes, we identify two distinct classes of autophagy foci. The first class are rapidly moving pre-phagophores, which are likely ATG9 vesicles that have begun to recruit autophagy factors. The second class of foci is phagophores that move more slowly as a result of ATG2mediated tethering of ATG9 vesicles to donor membranes. In addition, we show that most autophagy foci do not progress to become autophagosome marker positive. Intriguingly, our results suggest that ULK1 and ATG2 are maintained at low levels to prevent uncontrolled autophagosome production and are key signals in committing the phagophore into developing into an autophagosome. In total, our work provides critical mechanistic insight into autophagosome formation and establishes a new experimental framework for the quantitative analysis of autophagy in human cells.

## Results

### Genomic Insertion of HaloTag at Endogenous Loci of Autophagy Proteins

A common challenge in studying autophagy is the artificial structures formed when overexpressing autophagy proteins (Banerjee et al., 2020). Many overexpression artifacts have been successfully mitigated by stable expression of autophagy factors, for example by retroviral transduction, but multiple studies continue to confirm that this can still cause aberrant protein behavior (Banerjee et al., 2020; Barth et al., 2010; Kuma et al., 2007). To overcome this issue and maintain endogenous expression as well as regulation we used a well-established CRISPR genome editing strategy to introduce the HaloTag into the endogenous loci of autophagy factors that are involved in the initiation of autophagosome formation (ULK1, ATG13, PI4K3β), lipid transfer into the growing autophagosome (ATG2A and ATG9A), and LC3 conjugation (WIPI2, ATG16, ATG5, LC3) (Fig. 1A, Fig. S1A). Homozygous insertion of the HaloTag was confirmed by PCR and Sanger sequencing (Fig. S1B), and exclusive expression of the HaloTagged autophagy proteins was validated by western blot and fluorescent labeling (Fig. 1B,C). Due to their sequence complexity, we were unable to confirm homozygous editing of WIPI2 and ULK1 loci using PCR amplification. Instead, we validated the specificity of integration and confirmed that the cell lines exclusively expressed HaloTagged WIPI2 and ULK1 protein (Fig. S1C, Fig. 1B). To determine the expression levels of the HaloTagged autophagy proteins relative to the wildtype protein using western blots we lysed cells and removed the HaloTag using the TEV protease before gel electrophoresis. This approach avoids artifacts that we observed caused by the HaloTag affecting western blot transfer or antibody detection of the autophagy proteins (Fig. S2A-B). The majority of HaloTagged autophagy factors were expressed at similar levels to their untagged counterparts (Fig. S2A). HaloTagged ULK1 was approximately 4-fold overexpressed and WIPI2 expression was reduced 50% (Fig. S2B). Due to the sequence complexity of the WIPI2 locus, likely not all the WIPI2 alleles were modified causing a reduction in gene dosage. Importantly, HaloTagged-LC3 appeared to be more abundant than wildtype LC3 but the removal of the HaloTag by TEV cleavage eliminated the difference in the western blot signal (Fig. S2A-B). This demonstrates that the HaloTag can have a significant impact on the detection of proteins by western blot. This difference is likely caused by an alteration in western blot transfer efficiency since the non-specific binding of either the primary or secondary antibody to the HaloTag should have resulted in the appearance of an additional band corresponding to the HaloTag in samples treated with TEV, which was not observed (Fig. S2A). Altogether these observations demonstrate that we have successfully generated a collection of cell lines expressing HaloTagged autophagy proteins from their endogenous loci at or near the levels of their wildtype counterparts.

**Fig. 1.**
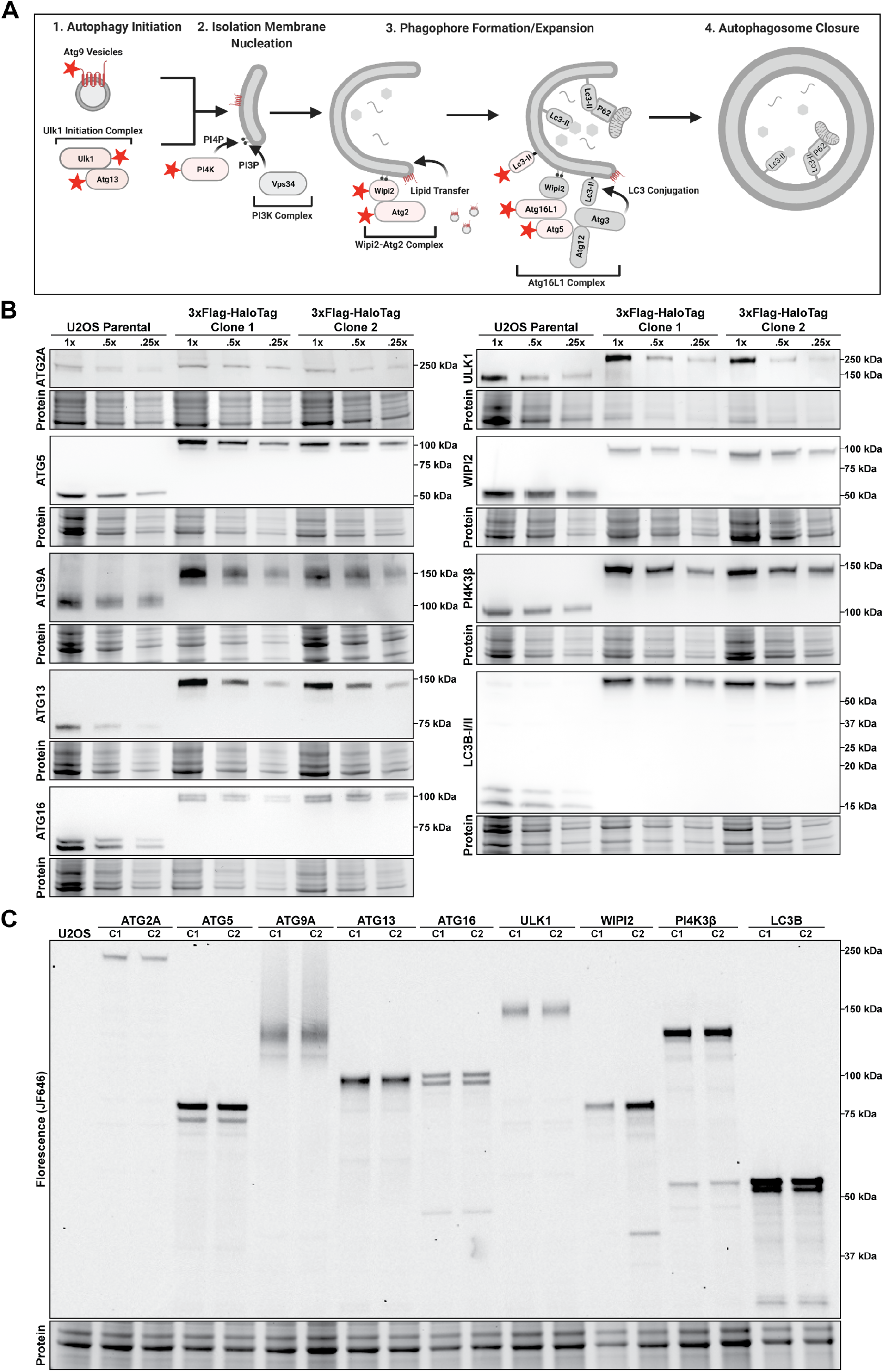
A HaloTag-based platform for quantitative analysis of autophagy in human cells. **(A)** Model showing autophagy complexes from the initiation membrane towards phagophore formation and autophagosome closure. Proteins tagged in this work are indicated with red stars. **(B)** Western blots of autophagy proteins showing size shift of the tagged protein and exclusive expression of the tagged protein in comparison to the parental U2OS cell line. Three concentrations (100, 50, and 25% of initial lysis volume) were loaded on the gel. **(C)** Fluorescence gel showing gene tagging; two distinct monoclonal lines (C1 and C2) were selected for each edited gene. Cell lines were labeled with saturating amounts of HaloTag ligand JF646 (250nM, 30min).

### The HaloTagged autophagy proteins are fully functional

Our quantitative western blots shown above demonstrate that our cell lines exclusively express HaloTagged autophagy proteins and allow us to evaluate the phenotypic impact of tagging the autophagy factors. To test the functionality of the HaloTagged autophagy proteins we determined the efficiency of autophagosome formation and degradation by measuring the levels of membrane conjugated LC3 after autophagy induction by rapamycin with and without the lysosome inhibitor bafilomycin, which prevents autophagosome degradation (Barth et al., 2010). LC3 western blots showed that apart from HaloTagged-ATG5, where we observed a minor conjugation defect, all cell lines expressing HaloTagged autophagy factors conjugated LC3 to a similar degree as the parental U2OS cells (Fig. S2C). We also carried out livecell imaging of autophagy proteins to verify the colocalization of tagged autophagy proteins with a well-established autophagosome marker. Cells were incubated with fluorescent HaloTag-ligand (JF646) alongside baculovirus-mediated transient expression of GFP-LC3. Surprisingly, GFP-LC3 did not form foci indicative of autophagosome formation, when combined N-terminal tags on the conjugation machinery including ATG5, ATG16, and WIPI2 (Fig. S3A). All the other tagged proteins formed puncta that colocalized with GFP-LC3 foci and responded similarly to the parental cell line when treated with rapamycin and bafilomycin (Fig. S3A).

Since we did not observe any defects in LC3 conjugation by western blot in HaloTag-ATG16 and HaloTag-WIPI2 cells, we hypothesized that the HaloTag sterically interferes with the conjugation of GFP-LC3. To test this hypothesis, we analyzed LC3 foci formation by immunofluorescence (IF) instead of GFP-tagged LC3. Using IF we detected similar numbers of LC3 foci within the Halo-ATG5, Halo-ATG16, and Halo-WIPI2 cell lines compared to parental U2OS cells, confirming that these HaloTagged autophagy factors can fully support endogenous LC3 conjugation (Fig. S3B). The formation of foci in response to autophagy induction is a characteristic of autophagy proteins. In contrast to LC3 foci number, which increased in response to rapamycin treatment (Fig. S3E), we did not observe increases in the number of foci formed by other HaloTagged autophagy proteins. HaloTagged ATG2, ATG13, ULK1, and LC3 foci number was only slightly increased in rapamycin-treated samples while HaloTagged ATG5, ATG9, and ATG16, showed no increase in foci formation (Fig. S3D). These observations suggested that inhibition of mTOR with rapamycin did not activate autophagy sufficiently to cause an increase in foci formation in our endogenously tagged cell lines. To induce autophagy using a more robust approach, we treated cells with Earl’s Balanced Salt Solution (EBSS), which triggers autophagy primarily by amino acid starvation. After treatment with EBSS, the number of foci formed by all HaloTagged autophagy factors was increased compared to cells grown in complete media (Fig. S3C). Strikingly, while the number of WIPI2 and ULK1 foci did not change after rapamycin treatment, amino acid starvation using EBSS lead to a 4-fold increase in the number of ULK1 and WIPI2 foci. When imaging HaloTag-PI4K we were unable to identify discreet foci but observed an accumulation of HaloTag-PI4K in the perinuclear region of the cytoplasm (Fig. S3C, F). This result differed from previous imaging experiments using stable expression cell lines to visualize PI4K. Notwithstanding, we observed a peri-nuclear redistribution of HaloTag-PI4K upon both rapamycin and amino acid starvation, which agrees with previous findings. In total, these results demonstrate that all of the tagged autophagy factors are functional and support autophagosome formation. Only HaloTag-ATG5 showed a minor defect in the rate of LC3 conjugation.

### Quantification of the absolute protein abundance of autophagy proteins

To determine the absolute number of protein molecules per cell for each tagged autophagy factor, we expanded upon an established in-gel fluorescence method to quantify the number of HaloTagged molecules per cell (Fig. 2A) (Cattoglio et al., 2019). To avoid the recombinant expression of each HaloTagged autophagy protein to use as a quantification standard, we supplemented U2OS cell lysates of a known number of cells with purified, recombinantly expressed 3xFLAG-HaloTag protein (Fig. 2B, S4A-E). Standard curves for the cell number using total protein levels and HaloTag-fluorescence signal were reproducible across all experiments with R2 values of 0.99 (Fig. S5A). These standard curves allowed us to precisely determine both the number of cells and the absolute number of HaloTagged molecules for each sample. Furthermore, we calculated a correction factor to compensate for fluorescence intensity differences caused by the banding pattern of the HaloTagged autophagy proteins by separating the HaloTag from the tagged protein using TEV protease (Fig. S2A-B). Finally, we added a correction factor to account for the relative expression of the HaloTagged protein compared to its wildtype counterpart (Fig. S5B-C). The in-gel fluorescence method allowed the quantification of autophagy protein abundance in U2OS cells over a broad range, from low expressed factors (ULK1, ~3,000 proteins/cell; ATG2, ~8,000 proteins/cell), to the highly expressed LC3 (>100,000 proteins/cell). ATG16 and ATG5, which form a complex, were present in a 1:1 ratio, consistent with their constitutive association. WIPI2, a scaffold protein that recruits ATG16-12-5 to the initiation membrane, was present in 5fold excess compared to ATG16. Surprisingly, ATG13 exceeded the abundance of ULK1 9-fold (Fig. 2C), considering that both proteins are part of a larger kinase complex (the ULK1 complex) (Mizushima, 2010). As an orthogonal approach, we determined the relative abundance of HaloTagged proteins in our collection of cell lines using flow cytometry and converted the relative fluorescence into an absolute protein number using HaloTag-ATG9 C1 as a fiducial point. The absolute protein abundance determined with flow cytometry and in-gel fluorescence were in good agreement, with some differences in the range of the technical error (Fig. 2D). Collectively, we have determined the absolute protein abundance of key autophagy factors and identified unexpected ratios between the subunits of the ULK1 complex. ULK1 and ATG2 are expressed at substantially lower levels than other autophagy proteins and may be key regulatory steps that throttle autophagic flux.

**Fig. 2.**
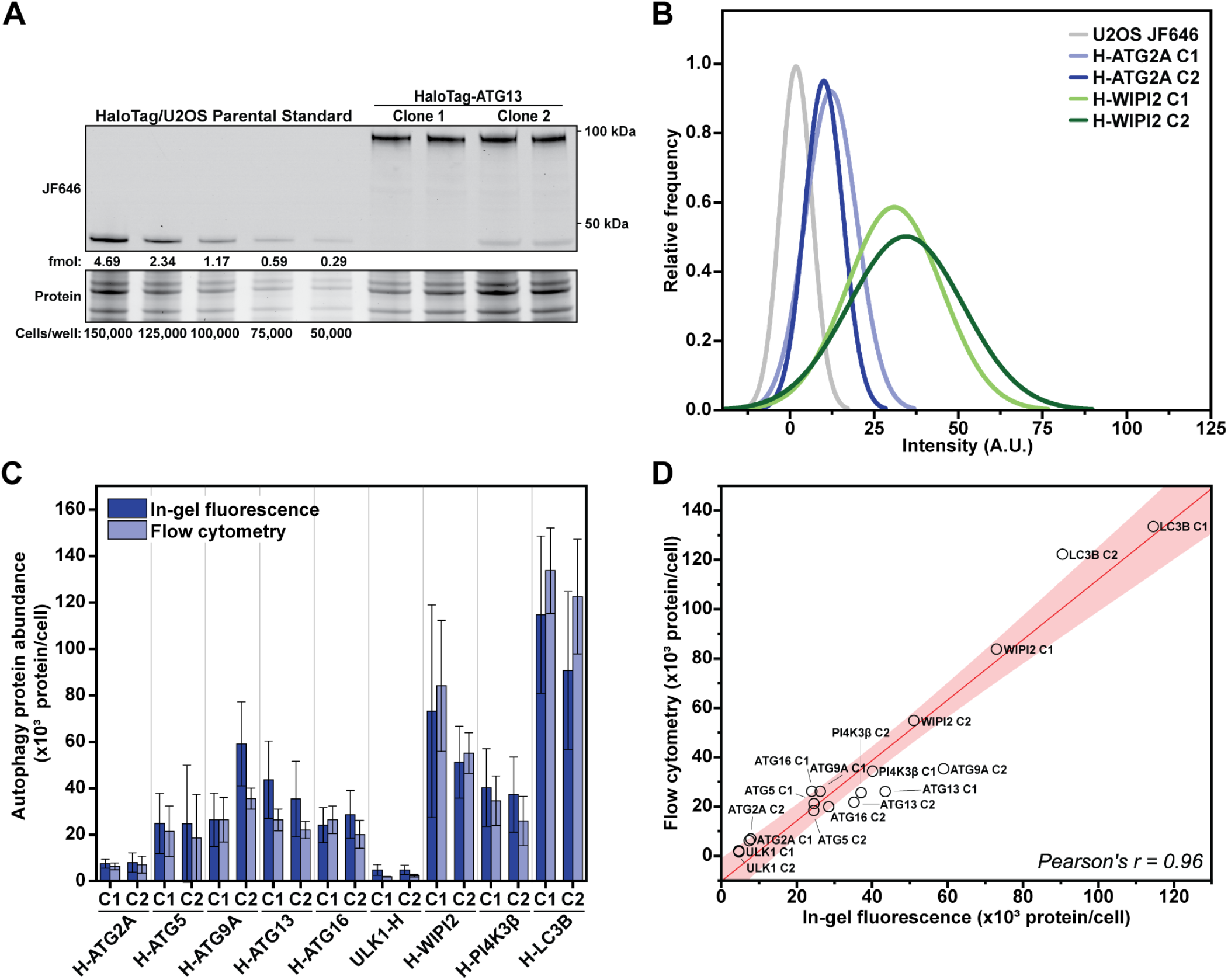
Absolute protein abundance quantification of autophagy factors in human cells. **(A)** Example in-gel fluorescence containing the quantification standards (HaloTag + cell lysate) and ATG13 protein. **(B)** Histogram of flow cytometry measurements depicting the relative protein abundances of U2OS (negative control), and two clones of cells expressing Halo-ATG2A, and Halo-WIPI2. **(C)** Corrected Protein abundance quantification of the tagged autophagy proteins with in-gel fluorescence and flow cytometry (N = 3, Mean ± SD including error propagation). **(D)** Graph showing the correlation between protein abundance measured by flow cytometry compared to in-gel fluorescence.

### Single-molecule analysis of the subcellular dynamics of autophagy factors

The dynamic recruitment of autophagy proteins to the sites of autophagosome formation is critical to controlling overall autophagic flux. Autophagosome formation can initiate at cargo-dependent sites such as mitophagy (Dalle Pezze et al., 2021). Alternatively, under starvation conditions, autophagosomes non-specifically engulf cellular material (Lamb et al., 2013). In either case, it is not known how autophagy factors encounter sites of autophagosome formation. Potential mechanisms include 3D-diffusion or scanning of existing membrane structures. To define the subcellular distribution of autophagy proteins and the mechanism by which they are recruited to the sites of autophagosome formation, we performed single-molecule live-cell imaging and single-particle tracking (SPT) of the

HaloTagged autophagy factors. Analysis of single-particle trajectories using the SpotOn tool allowed us to define distinct mobility states for each autophagy protein imaged (Fig. 3A-B, Movie S1-9). In principle, autophagy proteins could exist in three different states: freely diffusing in the cytoplasm, scanning a membrane (through intrinsic phospholipid binding, binding to a protein with intrinsic phospholipid binding, or interacting with transmembrane proteins), or statically bound to a site of autophagosome formation. To approximate the expected diffusion parameters for each of these states, we analyzed a HaloTag protein fused to a nuclear export signal (Halo-NES, Movie S10), to model freely diffusing proteins, and SNAP-SEC61, as a model membranebound protein (Movie S10). The HaloTag-NES diffused rapidly (D_free_ = 15 µm^2^/s, F_free_ = 99%) and had a negligible static fraction (F_Static_ = 1%) (Fig. 3C). SEC61 particles either slowly diffused (D_slow_ = 0.75 µm^2^/s, Fslow = 65%), or were static (D_static_ = 0.08 µm^2^/s, F_static_ = 35%) (Fig. 3C), which likely represents Sec61 molecules moving freely within the ER membrane or Sec61 molecules that are part of a translocon actively engaged with a ribosome in the process of translation, respectively.

**Fig. 3.**
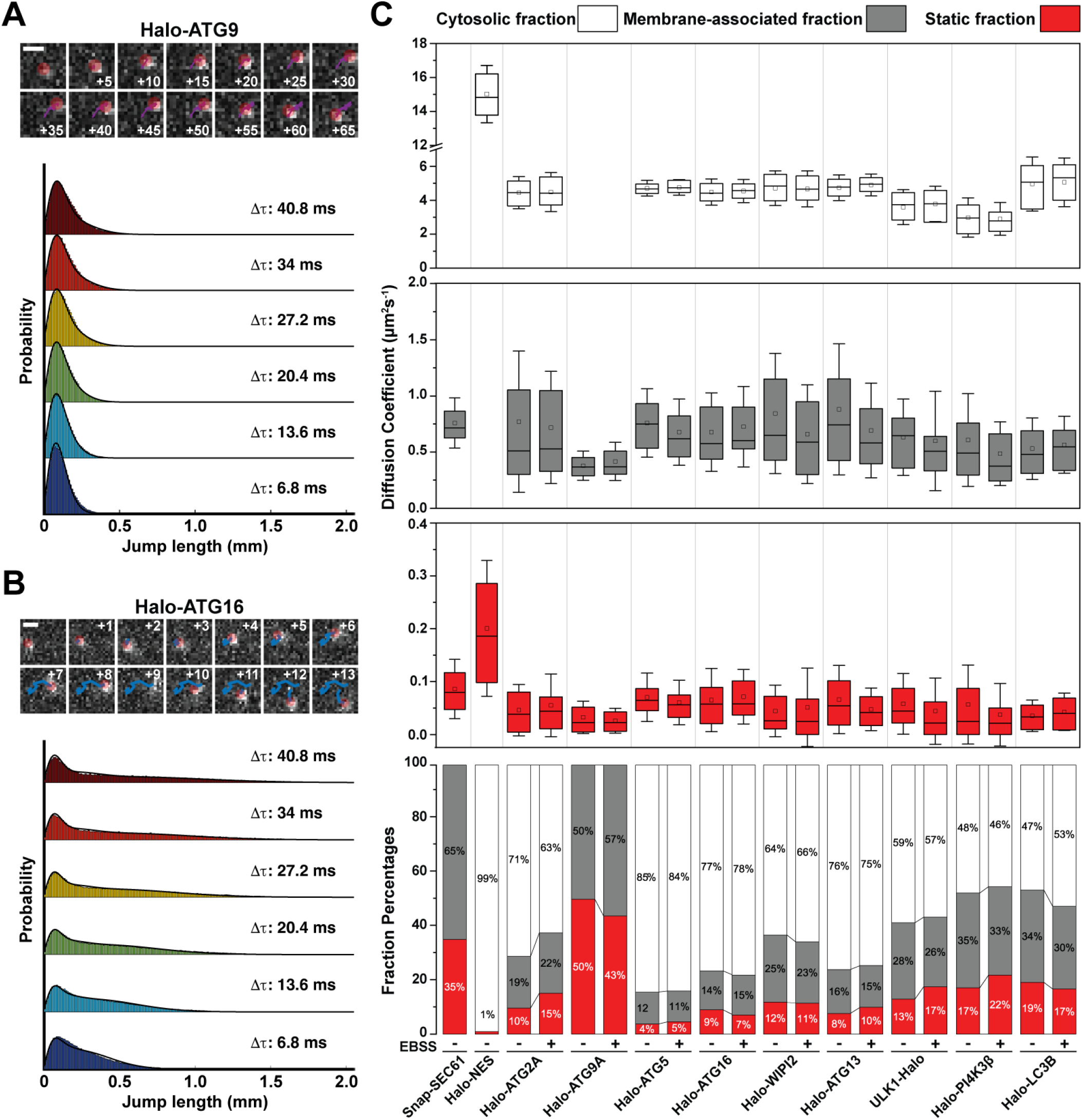
Live-cell single-molecule analysis of autophagy proteins. **(A-B)** Example of single-particle tracking of **(A)** ATG9 and **(B)** ATG16, and the corresponding fitting of the step-size probability distribution with SpotON algorithm. Numbers inside the micrographs indicate the imaging frame associated with the track. Movies were acquired at 6.8 ms per frame, scale bar = 1 µm. **(C)** Results of diffusive analysis for the HaloTagged autophagy proteins under control and EBSS starvation. Top 3 panels present the diffusion coefficients of the tracks based on the SpotON analysis. Bottom panel depicts the percentage associated with each fraction. The box indicates confidence interval ± SD, the square indicates the average, and the horizontal line is the median; for each condition, 3 biological replicates were analyzed, *∼*20 cells/replicate.

To define the diffusion properties of autophagy proteins, cells were imaged under control (control) and starved (EBSS treated) conditions. Except for ATG9, we used the 3-state model described above (freely diffusing, membrane scanning, statically bound to an autophagosome or other membrane) to fit the step size distributions of all autophagy proteins (Fig. 3B, S6). Since ATG9, like SEC61, is a trans-membrane protein and resides in lipid vesicles, it does not freely diffuse through the cytoplasm. Therefore, we applied a 2-state model to fit the step size distribution of ATG9 trajectories. The step-size distribution for all autophagy factors fits well with a 3-state model (or 2-state model in the case of ATG9, Fig. 3A, S6). For all other autophagy factors analyzed, a large fraction of the particles was freely diffusing (D_free_ = 4-6 µm^2^/s, F_free_ = 46-85%) (Fig. 3C). In addition, a significant fraction of molecules for all proteins analyzed moved with a diffusion coefficient comparable to SEC61 (D_slow_ = 0.5-0.75 µm^2^/s, F_slow_ = 12-35(Fig. 3C), consistent with these particles representing a membrane-associated population of the autophagy factors. Finally, a small fraction of molecules of all factors imaged were static (D_static_ = 0.02-0.08 µm^2^/s, F_static_ = 4-22%) (Fig. 3C). Approximately half of the ATG9 molecules diffused slowly (D_slow_ = 0.3 µm^2^/s, F_slow_ = 50%), while the other half of ATG9 particles were static (D_static_ = 0.02 µm^2^/s, F_static_ = 50%) (Fig. 3C). The diffusion properties of ATG5 and ATG16 were comparable, consistent with ATG5 and ATG16L1 forming a constitutive complex (Fig. 3C). Interestingly, the diffusion properties of ATG13 and ULK1 were distinct (Fig. 3C). ATG13 had a higher fraction of freely diffusion molecules than ULK1 (76% vs 59%) and the diffusion coefficient for this fraction was also higher for ATG13 compared to ULK1 (4.7 µm^2^/s vs. 3.6 µm^2^/s, p<0.001) (Fig. 3C). Together, this suggests that a substantial fraction of ATG13 is not associated with ULK1, which is consistent with our observation that the abundance of ATG13 exceeds the amount of ULK1 by approximately 9-fold. Freely diffusing PI4KIIIβ particles displayed the slowest diffusion coefficient of all proteins analyzed, which could be the consequence of transient interactions formed with endosomes and other Golgi-derived organelles (Judith et al., 2019; Waugh, 2019). Finally, most of the LC3 particles were in the bound or static state. LC3 exists in two primary forms, a lipid conjugated form (LC3-II) inserted into autophagic membranes and a cytosolic non-conjugated form (LC3-I) (Kabeya et al., 2004). Strikingly, the diffusion dynamics of none of the autophagy factors studied significantly changed after exposing cells to starvation conditions (Fig. 3C). This demonstrates that the dynamic properties of none of the factors studied are globally changed by cell starvation. In addition, the observation that the static populations remain unchanged under starvation conditions shows that only a very small fraction of molecules of a given autophagy factor are actively involved in autophagosome formation. Taken together, our single-molecule analysis of the diffusion dynamics of the tagged autophagy factors suggests that all proteins analyzed exist in a freely diffusing and membrane-associated state. In addition, our data demonstrate that the diffusion properties of the tagged autophagy proteins do not globally change in starvation conditions, indicating that only a small fraction of these proteins actively participates in autophagosome formation.

### Analysis of the autophagosome formation rate reveals two distinct populations of autophagosomes

We next sought to quantitatively analyze the recruitment of the tagged autophagy factors to autophagosomes, which was not possible using the single-molecule approach described above. The local accumulation of autophagy proteins at sites of autophagosome formation can be visualized as bright cytoplasmic foci and overall autophagic flux can be measured by determining the rate of foci formation using time-lapse microscopy (Dalle Pezze et al., 2021). Autophagy protein foci were automatically identified and tracked using a single particle tracking method (Fig. 4A,B; Movie S11-18) (Kuhn et al., 2021). Under control conditions, ULK1, ATG13, ATG5, and ATG16 formed significantly more foci (30-50 foci per cell per hour), compared to ATG2 and WIPI2, which formed a limited number of foci in control cells (~9 foci per cell per hour, Fig. 4C-D). Nutrient starvation significantly increased the number of foci formed by all autophagy factors imaged (Fig. 4C-D). Interestingly, the number of foci formed per cell per hour under starvation conditions was comparable (120-150) for ATG5, ATG13, ATG16, and ULK1, which was approximately 2-fold higher than the number of foci formed by ATG2 (~60 foci per cell per hour) (Fig. 4C-D). WIPI2 formed an intermediate number of foci (~90 foci per cell per hour) (Fig. 4C-D). This suggests that ATG2 is only detectably recruited to a subset of autophagosomes. To further analyze the characteristics of the autophagosomes detected, we determined the diffusion coefficient of cytoplasmic foci formed by the tagged autophagy factors, which reports on the mobility of the autophagosomes they associate with. The diffusion coefficient distribution of ATG2 positive autophagosomes revealed a single population with a mean diffusion coefficient of D = 0.002 µm^2^/s (Fig. 4E). In contrast, the diffusion coefficient distributions of all other autophagy factors analyzed were clearly made up of two distinct populations, one with a diffusion coefficient comparable to ATG2 positive autophagosomes, and a second population of foci that moved more rapidly (D = 0.01 µm^2^/s) (Fig. 4E). Importantly, the number of foci formed by ATG5, ATG13, ATG16, ULK1, and WIPI2 (e.g. F_slow_,ATG5 = 0.64 * 120 foci per cell per hour = 76 foci per cell per hour) that had a comparable diffusion coefficient to ATG2 foci is in a similar range as the number of ATG2 foci formed (~60 per cell per hour) (Fig. 4F).

**Fig. 4.**
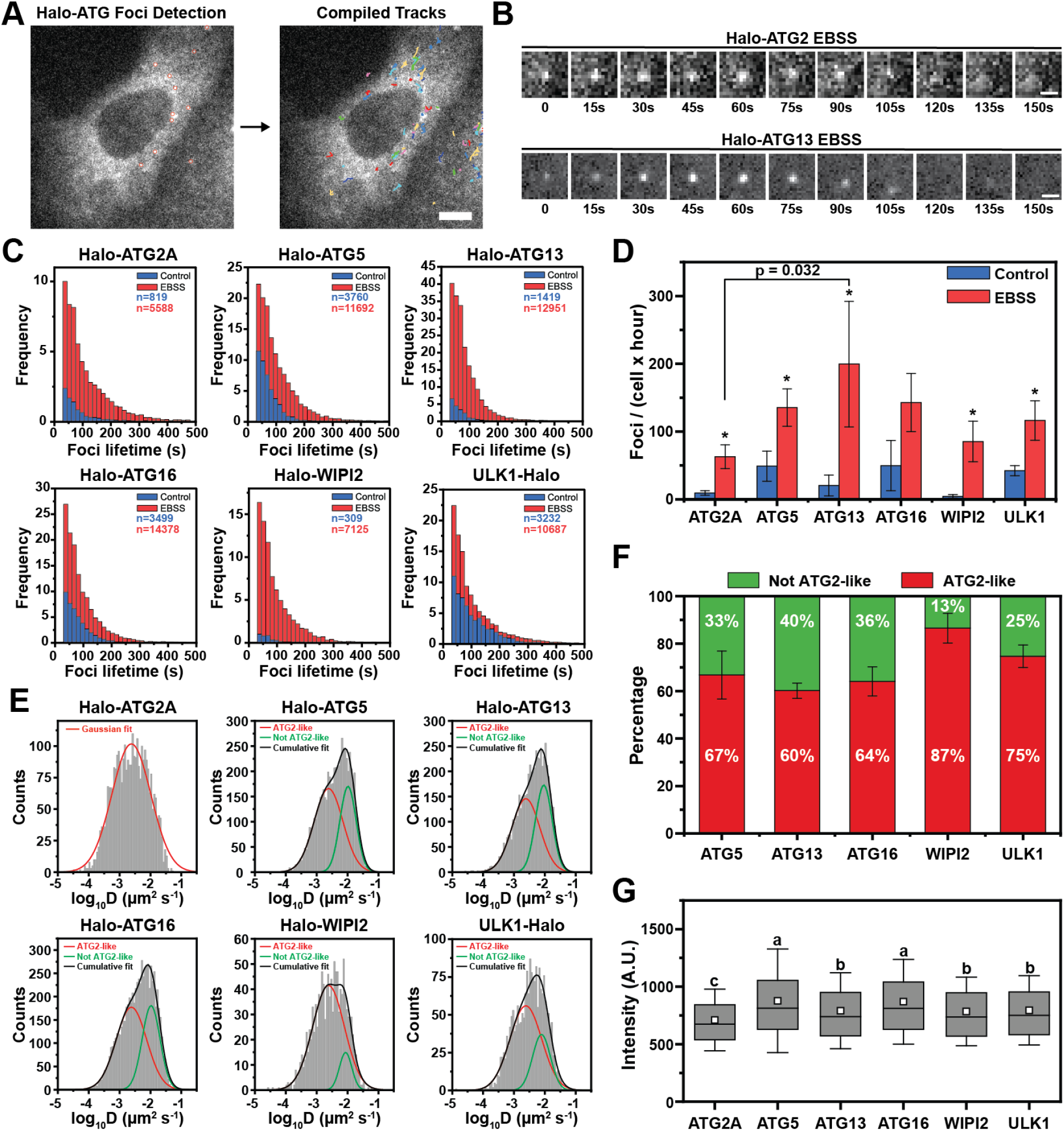
High-throughput quantification of autophagy foci lifetime and diffusion dynamics. **(A)** Upon labeling with fluorescent dye (JF646), cells expressing HaloTagged autophagy factors were starved (EBSS) and imaged at 4 frames per minute for 1h. TrackIT was used to detect foci based on threshold intensity (left) and connected into tracks using the nearest neighbor algorithm (right). **(B)** Example images of foci for Halo-ATG2A (upper panel) and Halo-ATG13 (bottom panel). Scale bar = 5 µm. **(C)** Histograms of foci lifetime for the HaloTagged autophagy proteins in control conditions (Control) and after 1h nutrient starvation (EBSS). Three biological replicates (20-30 cells per replicate) were performed for each HaloTag cell line. **(D)** Quantification of the number of foci formed per cell by autophagy factors over the course of 1h imaging in control (Control) and nutrient starvation (EBSS) conditions (N = 3, mean ± SD). A two-tailed t-test was used for statistical analysis (*p<0.05). **(E)** Histograms of diffusion coefficients of the foci formed by autophagy factors under nutrient starvation. Histograms were fitted with Gaussian curves. For the proteins other than Halo-ATG2, we fixed the mean of one subpopulation (in red) to match Halo-ATG2 mean. An additional subpopulation (in green) represent non ATG2-like foci. The black line represents the cumulative fitting. **(F)** Distribution of ATG2-like and non ATG2-like foci diffusion coefficients (N = 3, mean ± SD). **(G)** Quantification of the background corrected maximal intensity of the foci formed by the tagged autophagy factors. Boxes represent confidence interval ± SD, white square indicates the mean, horizontal line the median. Letters indicate statistically homogenous groups established by ANOVA (p < 0.05).

Finally, we determined the relative enrichment of the HaloTagged autophagy factors at sites of autophagy formation. Since all autophagy factors were fused to the same tag, and cells were labeled and imaged using identical conditions the peak intensity of the cytoplasmic foci detected is proportional to the number of protein molecules recruited to the autophagosome. ATG2 was the least abundant protein at sites of autophagosome formation, while ULK1, ATG13, WIPI2, ATG5, and ATG16 were slightly more abundant than ATG2 (Fig. 4G). The mean peak intensity of ULK1 and ATG13 as well as ATG5 and ATG16 were comparable, indicating that similar amounts of these proteins are recruited to autophagosomes, consistent with them being present on autophagosomes as part of their known protein complexes (Fig. 4G). Together these observations suggest that ATG5, ATG13, ATG16, and ULK1 are recruited to all autophagosomes while ATG2 only detectably accumulates at around half of the autophagy foci. In addition, our analysis of the diffusion coefficients of foci formed by the tagged autophagy factors suggests that the ATG2 recruitment triggers the transition of the phagophore to a less mobile state, potentially by the formation of multiple anchor points to lipid donor membrane compartments by ATG2.

### A positive feedback loop involving ULK1 complex and VPS34 is critical for the initiation of autophagosome formation

The data presented thus far demonstrate that our automated particle tracking approach can quantitatively determine the frequency and biochemical and physical properties of starvation foci induced formation by all tagged autophagy factors. To dissect the genetic requirements for the initiation of autophagy-induced foci by the tagged autophagy factors, we first focused on the ULK1 complex. It is well established that ULK1 is critical for the initiation of autophagy (Karanasios et al., 2016; Mercer et al., 2009; Mizushima, 2010; Szymańska et al., 2015). As a marker for the ULK1 complex, we used HaloTag-ATG13. HaloTag-ATG13 was clearly recruited to “ATG2-like” static autophagosomes and foci with higher mobility, which we will refer to as untethered pre-phagophores from here on. To dissect the contribution of the ULK1 complex to the formation of both classes of autophagosomes, we knocked out ULK1, FIP200, or ATG101 in the HaloTag-ATG13 cell line (Fig. 5A). Clonal knockout cell lines of ULK1, FIP200, or ATG101 showed a significant defect in LC3 conjugation under starvation conditions and a shifted phosphor-P62 band within the FIP200 and ATG101 knock-out (Fig. 5B). Interestingly, LC3 conjugation was more strongly inhibited in FIP200 and ATG101 knockout cells compared to cells lacking ULK1 (Fig. 5B,C). It is possible that ULK2, compensates for the loss of ULK1 activity. We were unable to generate ULK1 and ULK2 double knock-out cells, indicating that loss of ULK1 and ULK2 function may be lethal in U2OS cells. As an alternative approach, we combined the ULK1, FIP200, and ATG101 gene knockouts with ULK-101 a highly potent and specific small molecule inhibitor of both ULK1 and ULK2 (Martin et al., 2018). ULK-101 treatment in ULK1 knock-out cells further decreased LC3 conjugation, to the levels observed in cells lacking FIP200 or ATG101, confirming the previously reported role for ULK2 in maintaining LC3 conjugation in the absence of ULK1 (Fig. 5B,C). In addition, these observations demonstrate that knock-out of ATG101 and FIP200 lead to a complete loss of ULK1 complex function, which is consistent with previous studies (Itakura and Mizushima, 2010; Kannangara et al., 2021). To determine which step of autophagosome formation is inhibited by the loss of function of ULK1, FIP200, and ATG101, we assessed the formation of ATG13 foci in the knock-out cell lines in both control and starved conditions (Fig. 5D). As expected, under control conditions HaloTag-ATG13 formed a limited number of foci in all cell lines (Fig. 5E, Movie S19). After treatment with EBSS, HaloTag-ATG13 foci formation was dramatically to ~200 foci/cell/hour in the parental cells (Fig. 5E). Knockout of FIP200 and ATG101 resulted in a significant reduction in the number of HaloTag-ATG13 foci formed, from 200 to 9 and 15 foci/cell/hour, respectively (Fig. 5E, Movie S19).

**Fig. 5.**
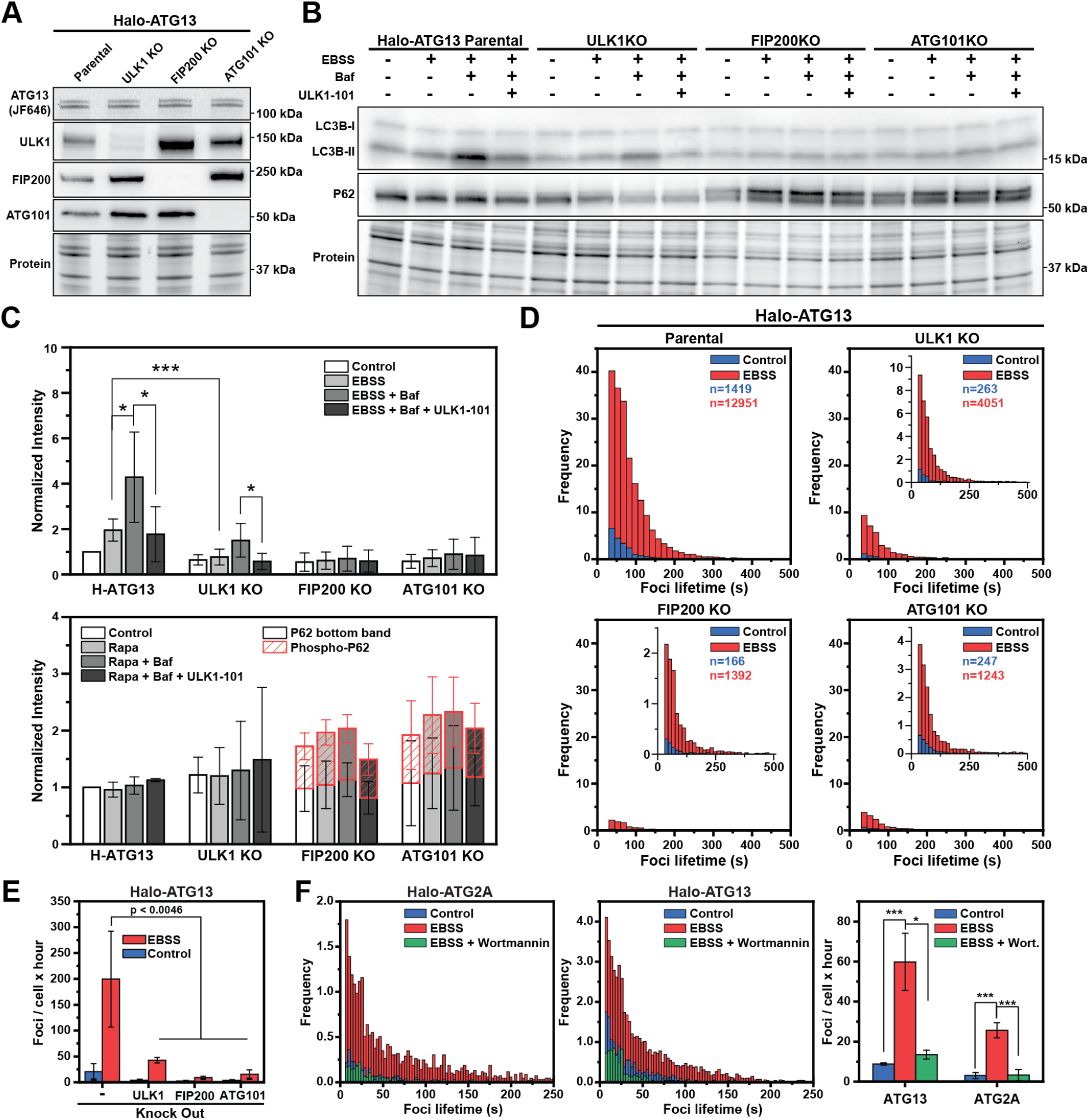
The Ulk1 complex is critical for autophagosomes initiation. **(A)** Fluorescence gel and western blots demonstrating successful knock-out of ULK1, FIP200 and ATG101 from the Halo-ATG13 cell line. **(B)** Western blots demonstrating impaired autophagy when ULK1, FIP200 and ATG101 are individually depleted from the Halo-ATG13 cell line. For the treatment experiments, cells were preincubated in control media with ULK1-101 (1µM) or without drug for 1 hour where indicated. Cell were then switched to their control or EBSS starvation media, with or without bafilomycin (100nM), for an additional hour. **(C)** Quantification of the western blots in (B). Data represent mean ± SD over three biological replicates. Phospho-P62 band (red striped bar graph) was detected and quantified only in the FIP200 and ATG101 knock-out cell lines. **(D)** Histograms of Halo-ATG13 foci lifetimes for the parental Halo-ATG13 cells and ULK1, FIP200, ATG101 knock-out cell lines in control conditions (Control) and after 1h nutrient starvation (EBSS). Three biological replicates (20-30 cells per replicate) were performed for each cell line. **(E)** Quantification of the number of foci formed per cell by HaloTag-ATG13 over the course of 1h imaging in control (Control) and nutrient starvation (EBSS) conditions (N = 3, mean ± SD). A two-tailed t-test was used for statistical analysis. **(F)** Histograms of foci lifetimes for the HaloTag-ATG2A (left) and HaloTag-ATG13 (center) in control and nutrient starvation (EBSS, 1h) conditions with and without wortmannin (1µM). Cells were pretreated with wortmannin for 1 hour. Right panel presents the quantification of the foci frequency. Data represent mean ± SD over three biological replicates (20-30 cells per replicate). A two-tailed t-test was used for statistical analysis (*p <0.05, **p <0.01, ***p <0.001).

ULK1 knock-out had an intermediate phenotype with ~50 foci/cell/hour (Fig. 5E), consistent with the intermediate phenotype observed for LC3 conjugation. Together these observations suggest that the ULK1 complex is required for all autophagosomes. A critical consequence of ULK1 activation is thought to be the localized generation of PI3P by VPS34 at the site of autophagosome formation. To determine if PI3P accumulation is clearly downstream of ULK1 complex enrichment at the phagophore, we treated HaloTag-ATG13 cells with 1 µM Wortmannin, which inhibits the PI3kinase activity of VPS34, for 1 hour prior to and during cell starvation with EBSS and analyzed foci formation by HaloTag-ATG13. Strikingly, inhibition of PI3P formation reduced the number of HaloTag-ATG13 foci formed under starvation conditions to levels comparable with cells that are grown in complete media (Fig. 5F, Movie S20). This suggests that PI3P formation is critical for ATG13 accumulation at the phagophore. As a control, we treated HaloTag-ATG2 cells with wortmannin and analyzed its impact on HaloTag-ATG2 foci formation. Similar to HaloTag-ATG13, the number of HaloTag-ATG2 foci formed in starved cells was reduced to those observed under control conditions by treatment with wortmannin (Fig. 5F, Movie S21). Importantly, this experiment confirmed that the number of HaloTag-ATG2 foci formed is approximately half of the number of HaloTag-ATG13 foci formed in all experimental conditions (Fig. 5F). Since these experiments were carried out at a higher time resolution compared to the experiments shown in Figure 4, we also confirmed that the diffusion coefficient distribution of ATG13 foci contained two distinct populations, while that of ATG2 only had a single slowly diffusing population (Fig. S7C). These observations demonstrate that PI3P formation at the phagophore is required for detectable accumulation of ATG13 and likely the whole ULK1 complex at sites of autophagosome formation, consistent with previous observations by others (Karanasios et al., 2013a). We hypothesize that a positive feedback loop initiated by ULK1 activation and reinforced by additional ULK1 complex recruitment via the formation of PI3P by VPS34 at the phagophore is the critical trigger for autophagosome formation. Importantly, ATG13 contains a PI3P/PI4P binding sequence in its N-terminus, which is thought to contribute to the association of the ULK1 complex with the phagophore (Karanasios et al., 2013a). To analyze the dynamic association of ATG13 with cytoplasmic membranes we took advantage of the live-cell single-molecule imaging approach we developed. Our observations demonstrated that a fraction of HaloTag-ATG13 particles were highly static and a second subset of HaloTag-ATG13 molecules moved with a similar diffusion coefficient as SNAP-Sec61. We propose that these slowly moving and static HaloTag-ATG13 particles are the consequence of the association of HaloTag-ATG13 with cytoplasmic membrane compartments or the phagophore. We first determined whether the diffusion properties of HaloTag-ATG13 were impacted by the knock-out of ULK1, FIP200, or ATG101. HaloTag-ATG13 diffusion properties were unchanged in cells lacking ULK1, FIP200, or ATG101 under both control and starved conditions (Fig. S7A-B, Movie S22). These observations suggest that the membrane association of ATG13 does not depend on its association with any of the other ULK1 complex components. Our results demonstrate that the initiation of autophagosome formation requires the ULK1 complex. In addition, ULK1 complex accumulation at the phagophore requires PI3P, which is potentially mediated by the association of ATG13 with PI3P. Altogether, our observations are consistent with a model in which a positive feedback loop initiated by the ULK1 complex and amplified by VPS34 is critical to trigger autophagosome formation.

### The majority of cytoplasmic foci formed by autophagy proteins do not progress to form mature autophagosomes

The results described thus far were focused on the initiation of starvation-induced foci by the tagged autophagy proteins. To analyze the progression of these foci into mature autophagosomes, we performed quantitative dual-color live-cell imaging to assess the recruitment of P62 and LC3 to the starvation-induced foci formed by the tagged autophagy factors. Cells expressing the HaloTagged autophagy proteins were transduced with baculoviruses encoding GFP-LC3 or GFP-P62 and imaged with high time resolution (3 s per frame) after cell starvation with EBSS (Fig. 6A, Fig. S8A,Movie S23-32). GFP and HaloTag signals were tracked independently, and co-localization was defined as particles whose centroids were less than 3 pixels apart at any given timepoint (see methods for details). This approach allowed us to analyze hundreds of autophagosomes in a completely unbiased fashion. We analyzed all tagged proteins except for PI4K and ATG9, which did not form foci. A fraction of all autophagy factors analyzed co-localized with P62 (P62_positive_, Fig. 6C),but the majority of foci never showed detectable P62 accumulation (P62_negative_, Fig. 6C). Importantly, the overall rate and lifetime of P62 foci was comparable across all cell lines, indicating that the HaloTagged autophagy proteins do not impact the formation of P62-positive autophagosomes (Fig. 6D). Importantly, the fraction of HaloTag-ATG2 foci that colocalized with P62 (23%) was approximately double that of all other autophagy proteins tested (10%), which suggests that ATG2 recruitment is a critical step in the commitment of the phagophore to mature into an autophagosome (Fig. 6F).

To gain further insight into the assembly order and timeframe of autophagosome formation we analyzed the kinetics of foci formation and co-localization with P62. The lifetime of the P62_negative_ trajectories (Mean = 23-41 s) was significantly shorter than the P62_positive_ trajectories (Mean = 91-122s) for all tagged autophagy proteins (p<0.001) (Fig. 6E). On average autophagy protein foci formed 40 seconds prior to the recruitment of P62, co-colocalized with P62 for 20 seconds, and dissociated from P62 foci 30 seconds prior to the disappearance of the P62 signal, which is likely the consequence of autophagosome fusion with the lysosome and quenching of the GFP signal (Fig. 6B). The total time of autophagosome formation (ATG signal appearance to P62 signal disappearance) was similar for all proteins tested (150 seconds) (Fig. 6B). Importantly, we confirmed these results using GFP-LC3 for the HaloTagged cells lines that support GFP-LC3 foci formation (ATG2, ATG13, ULK1, see above) (Fig. S8B). Altogether, these results demonstrate that most autophagosome initiation sites rapidly disassemble after their formation and do not progress to form mature autophagosomes. This raises the question of which molecular events are critical to driving the transformation of a phagophore into an autophagosome. Our observations indicate that the recruitment of ATG2 is one of the key steps in the maturation process of the autophagosome. Importantly, if the phagophore matures into an autophagosome all autophagy factors analyzed are recruited in close succession approximately 40 seconds prior to the accumulation of P62. Finally, our results demonstrate that the average lifetime of an autophagosome is 150 seconds.

**Fig. 6.**
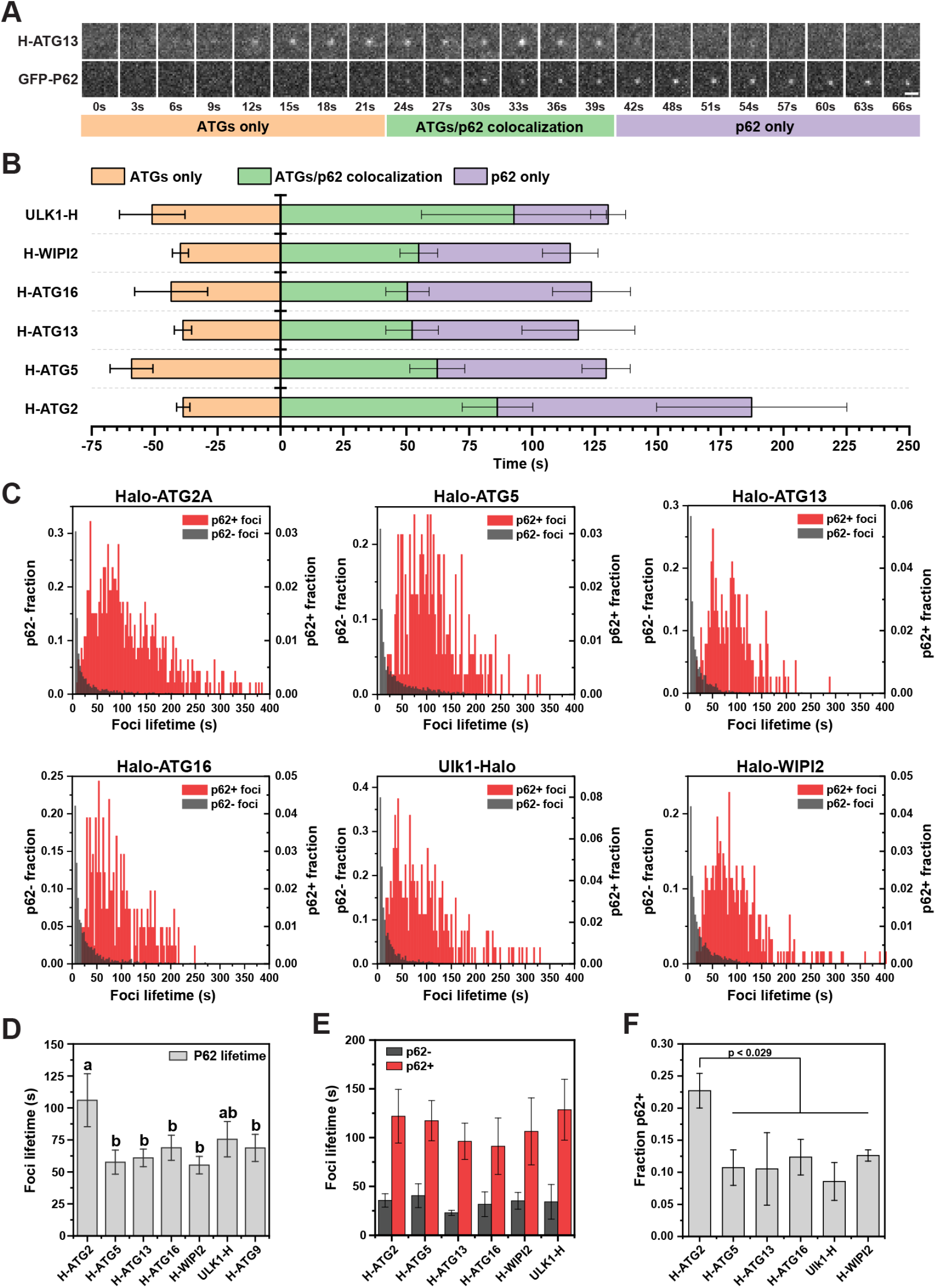
Analysis of the maturation kinetics of autophagosomes using dual-color imaging. **(A)** Example images showing formation, growth and disappearance of colocalized Halo-ATG13 (top) and GFP-p62 foci (bottom) using dual-color live-cell imaging under EBSS starvation (1h). Scale bar = 2 µm. **(B)** Quantification of the timing of the three distinct phases of autophagosome formation for the HaloTag proteins under EBSS starvation (1h). Data represent mean ± SD of three biological replicates (20-30 cells per replicate). **(C)** Histograms of the lifetime of HaloTagged autophagy protein foci that colocalized (red) or did not colocalize (dark grey) with GFP-P62. **(D)** Quantification of the lifetime of GFP-P62 foci in the HaloTagged autophagy factor cell lines. Data represent mean ± SD of three biological replicates (20-30 cells per replicate). Letters indicate statistically homogenous groups established by ANOVA (p < 0.05). **(E)** Quantification of the average HaloTagged autophagy protein foci lifetime for GFP-P62 positive (red) and negative (dark grey) foci. Data represent mean ± SD of three biological replicates (20-30 cells per replicate). **(F)** Percentage of HaloTagged autophagy protein foci that co-localized with P62 foci. Data represent mean ± SD of three biological replicates (20-30 cells per replicate). A two-tailed t-test was used for statistical analysis.

### ATG9 does not detectable accumulate at sites of autophagosome formation

A growing body of evidence suggests that ATG9-containing vesicles are the seed upon which the phagophore is assembled (Sawa-Makarska et al., 2020). To test this model, we knocked out ATG9A in the HaloTag-ATG2 cell line (Fig. 7A). Compared to the parental H-ATG13 cell line, the ATG9A knock-out shows a molecular size shift on the P62 band attributable to phospho-P62 (Fig. 7A), which is an indication of impaired autophagy. We then analyzed foci formation of HaloTag-ATG2 in the ATG9A knock-out (Fig. 7A). Strikingly, HaloTag-ATG2 foci formation was largely eliminated in the absence of ATG9A under both control and starvation conditions (Fig. 7B-D, Movie S33). The limited number of foci formed by HaloTag-ATG2 in the absence of ATG9A could be triggered by ATG9B, a lowly expressed isoform (Kusama et al., 2009). Attempts to remove ATG9B from ATG9A knock-cells by knock-out or knock-down were unsuccessful, indicating that ATG9B is essential in the absence of ATG9A. The reduction in HaloTag-ATG2 foci observed in the absence of ATG9A suggests that ATG9A-containing vesicles constitute the platform for phagophore formation or ATG9A activity is required to grow autophagosomes to a sufficient size to facilitate the detectable accumulation of HaloTag-ATG2. Importantly, while P62 foci formation kinetics were identical in HaloTag-ATG2 and HaloTag-ATG9A cells (Fig. 6E), we failed to detect accumulations of HaloTag-ATG9A that co-localized with P62 foci. In contrast, as shown above (Fig. 6F), a large fraction of HaloTag-ATG2 foci co-localize with P62. This suggests that, while ATG9A is critically important for autophagosome formation, the number of ATG9A molecules that are recruited to autophagosomes is not detectable by our current method. These observations are consistent with ATG9 vesicles functioning as the seeds for autophagosome formation limiting the number of ATG9 molecules contained in the phagophore to those that were present in the original ATG9 vesicle.

**Fig. 7.**
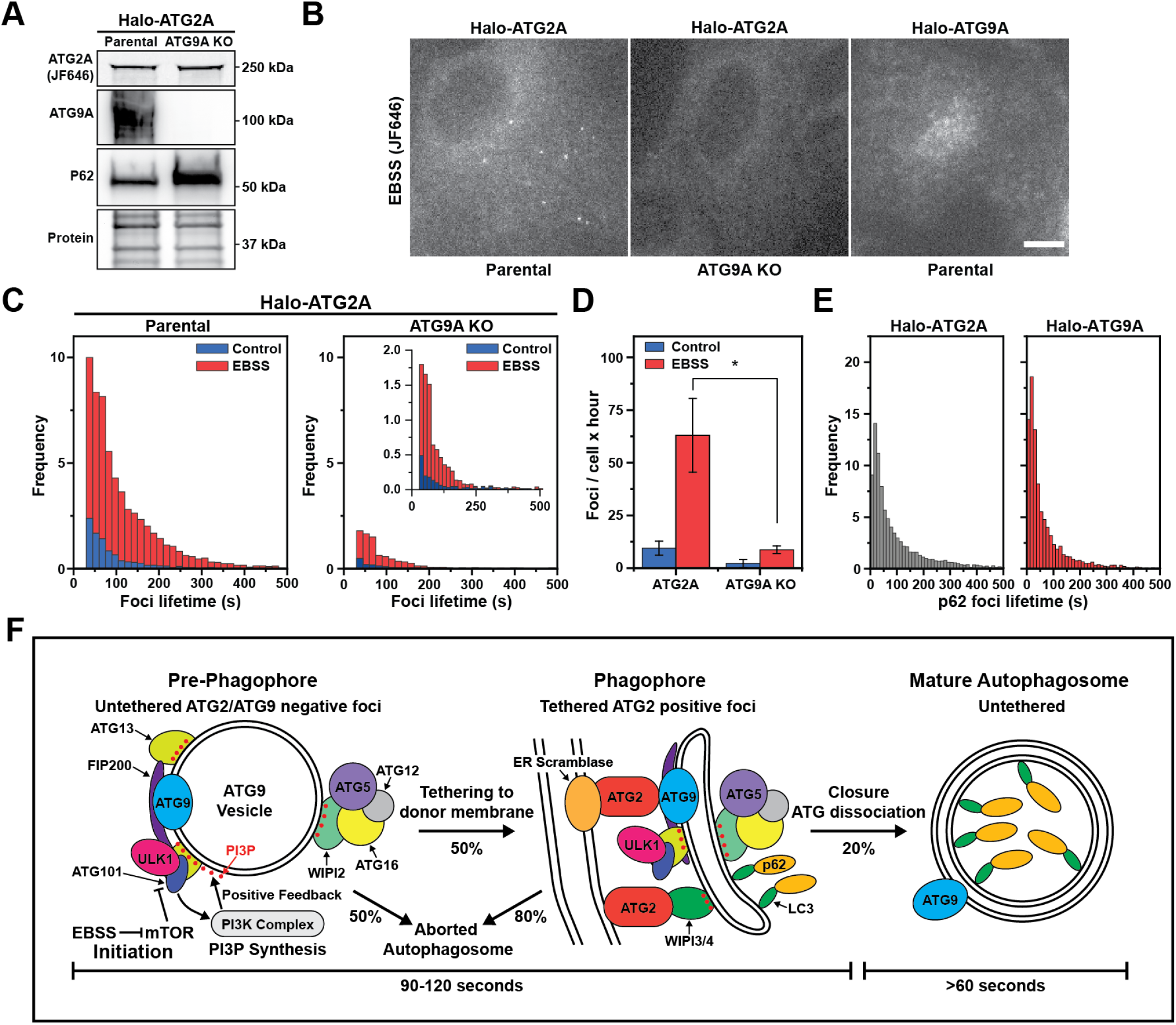
ATG9 vesicles constitute the platform for autophagosome formation. **(A)** Fluorescence gel and western blots demonstrating successful ATG9A gene knock-out from cells expressing HaloTag-ATG2A. The ATG9A knock-out cells accumulate P62, indicating impaired autophagy. **(B)** Micrographs showing decrease of Halo-ATG2A foci when ATG9A is depleted (central panel). Halo-ATG9A does not form foci detectable foci under EBSS starvation (right panel). **(C)** Histograms of HaloTag-ATG2 foci lifetime in parental and ATG9A knock-out cells (EBSS, 1h). **(D)** Quantification of the number of foci formed per cell over the course of an hour by HaloTag-ATG2 in control and ATG9A knock-out cells. Data represent mean ± SD of three biological replicates (20-30 cells per replicate). A two-tailed t-test was used for statistical analysis (*p<0.05). **(E)** Histograms of GFP-P62 foci lifetime in cells expressing HaloTag-ATG2A (left) and HaloTag-ATG9A (right) under nutrient starvation (EBSS, 1h). **(F)** Model for the formation of the autophagosome. Upon mTOR inhibiton, a ULK1 complex - PI3K feedback loop initiates the assembly of autophagy proteins on an untethered ATG9 vesicle. Half of these pre-phagophore structures are tethered to cellular membranes via ATG2A. Only 20% of the ATG2A positive phagophores mature to a point where P62 is detectably recruited. The formation and growth of a full autophagosomes takes about 90-120 seconds. Upon closure, the autophagy proteins dissociates, the mature autophagosomes is unthethered and is delivered to lysosomes (>60s).

## Discussion

The experiments presented in this study systematically and quantitatively assess the formation of autophagosomes in human cells. Our work provides new mechanistic insights into the initiation and maturation of autophagosomes and the collection of cell lines we have created expressing HaloTagged autophagy factors from their endogenous loci represents a tremendously valuable tool for future studies of autophagosome formation in human cancer cells.

### ATG9 vesicles constitute the platform for autophagosome formation

ATG9A/B are the only known transmembrane proteins that are critical for autophagosome formation (Guardia et al., 2020). Several recent structural and biochemical studies have demonstrated that ATG9A forms a trimeric membrane pore that functions as a lipid scramblase, passively exchanging lipids between the inner and outer leaflets of lipid bilayers (Guardia et al., 2020; Matoba et al., 2020; Noda, 2021). These observations have led to a model in which ATG9-containing vesicles constitute the platform upon which the autophagosome is formed via ATG2-mediated lipid transfer from other membrane sources such as the endoplasmic reticulum (ER). Work by others has shown that ATG9 containing proteo-liposomes or immunopurified ATG9 vesicles are sufficient for reconstituted nucleation of the autophagosome (Sawa-Makarska et al., 2020). While these in vitro observations are consistent with the idea that ATG9 vesicles are the platform for autophagosome formation, the evidence that ATG9 vesicles transform into autophagosomes in cells is limited.

The introduction of a HaloTag at the endogenous ATG9A locus facilitated the highly sensitive detection of ATG9A in living cells. Our observations revealed that ATG9A is the only autophagy factor analyzed that does not form foci and therefore is not detectably enriched at sites of autophagosome formation. Importantly, we also demonstrate that ATG9A is essential for the accumulation of ATG2A at starvation-induced foci in U2OS cells. Together these results strongly support a model in which ATG9A containing vesicles are the platform for autophagosome formation. The number of ATG9A molecules contained in an autophagosome is limited to the molecules that were present in the ATG9 vesicle that was specified to initiate autophagosome formation. Additional ATG9A molecules could only be recruited to the phagophore by fusion with other ATG9Acontaining vesicles. While our experiments cannot completely exclude the possibility that multiple ATG9A vesicles fuse in the context of autophagosome maturation, our observations suggest that the total number of ATG9A molecules incorporated into the phagophore is significantly less than other autophagy factors. In particular, ATG2A, a proposed direct interactor of ATG9A (Gómez-Sánchez et al., 2018), forms robust foci at autophagosomes, potentially by WIPI3/4-mediated recruitment of ATG2 to the phagophore (Chowdhury et al., 2018; Otomo et al., 2018; Ren et al., 2020). Importantly, if ATG9-containing vesicles were transformed into phagophores and eventually autophagosomes, one would not expect to observe a substantial accumulation of ATG9A at sites of autophagosome formation. We, there-fore, believe that the results presented in this study provide direct evidence that ATG9 vesicles constitute the platform for autophagosome formation in human cancer cells (Fig. 7F).

### The ULK1 complex and PI3-kinase form a positive feedback loop required for autophagosomes formation

A key open question is, how ATG9 vesicles are specifically selected to transform into autophagosomes. The ULK1 complex, composed of ULK1, ATG13, FIP200, and ATG101 is essential in the initiation of autophagy (Dikic and Elazar, 2018; Lin and Hurley, 2016; Yu et al., 2018). Work by others has demonstrated that ATG13 and FIP200 are required for the localization of ULK1 to the phagophore (Chang et al., 2021a; Ganley et al., 2009; Shi et al., 2020). A key function of the ULK1 complex is the activation of the PI3-kinase VPS34 at sites of autophagosome formation to promote the production of PIP3 (Mercer et al., 2018; Park et al., 2016; Russell et al., 2013). In addition, it has been shown that ULK1 complex recruitment to the phagophore is re-enforced by the association of ATG13 with PIP3 (Karanasios et al., 2013a). Therefore, the ULK1 complex and PI3-kinase form a positive feedback loop required to initiate autophagosome formation (Ohashi, 2021).

The work presented in this study further confirms this model. The formation of starvation-induced foci by ATG13 was abolished when cells were pre-treated with wortmannin to inhibit PI3-kinase activity, demonstrating that PIP3 is essential for the detectable accumulation of ATG13 at sites of autophagosome formation. In addition, the diffusion dynamics of ATG13 were largely unchanged when the other components of the ULK1 complex were knocked out, which demonstrates that the ULK1 complex likely forms in the context of the phagophore and does not assemble into a preformed complex in the cytoplasm. Importantly, while knockout of FIP200 and ATG101 strongly reduced LC3 conjugation, ULK1 knock-out cells retained the ability to carry out a reduced amount of LC3 conjugation, which was further reduced by treatment with ULK101. This observation demonstrates that ULK1 and ULK2 both play critical roles in driving autophagosome formation in U2OS cells. Finally, ATG13 is 10-fold more abundant than ULK1 and given its ability to interact with PIP3 enriched membranes we postulate that ATG13 recruitment is one of the earliest events in autophagosome formation. In future work it will be critical to define the molecular mechanism by which the ULK1 complex and PI3kinase are directed to ATG9 vesicles to initiate autophagosome formation (Ren et al., 2022).

### ATG2 recruitment leads to phagophore immobilization

Once an ATG9 vesicle has been specified as a seed for autophagosome formation it has to expand, engulf cargo, and eventually close into a mature autophagosome. ATG2 is an extended barrel shaped protein that can form tethers between lipid vesicles and transfer phospholipids between them (Chowdhury et al., 2018; Noda, 2021; Tang et al., 2019; Valverde et al., 2019). Therefore it is thought that ATG2 is the critical factor for transferring phospho-lipids from a donor lipid source to the growing phagophore.

Our imaging methodologies allowed us to systematically analyze the recruitment of ATG2 to the phagophore. Strikingly, the number of ATG2 foci formed was approximately half of the number formed by all other autophagy factors tested except for WIPI2. In addition, the diffusion coefficient distribution of ATG2 showed a single slowly moving population, while all other factors contained two populations, one which was comparable to ATG2 positive foci and a second more rapidly diffusing population. Importantly, the number of slowly diffusing autophagy factor foci was similar to the number of ATG2 foci observed, suggesting that they reflect the same population of phagophores. These observations demonstrate that ATG2 marks a subset of phagophores with distinct biophysical properties. We propose that the reduced diffusion coefficient of the ATG2 positive phagophore population is the result of ATG2 mediated tethering of the phagophore to donor membranes. Importantly, these observations further support the model that ATG9 vesicles are the seed for autophagosome formation. The rapidly diffusing population of autophagy foci (D = 0.1 µm^2^/s) is comparable to the diffusion coefficient of ATG9 vesicles measured in our single-molecule imaging experiments (D = 0.2 µm^2^/s). Immobilization of an ATG9 vesicle by tethering it to other cellular membranes via ATG2 would be expected to reduce their diffusion coefficient, which is consistent with our observations. In contrast, it is not immediately apparent why ATG2 recruitment should reduce the mobility of the phagophore if it was formed by an alternative mechanism. In addition, our observations suggest that the ULK1 complex, PI3-kinase, the LC3 lipidation machinery, and to a smaller degree WIPI2 can be recruited to ATG9 vesicles prior to their tethering to donor membranes by ATG2.

Together these observations demonstrate that the recruitment of ATG2 marks the transition from an ATG9 vesicle into a phagophore that can expand and mature into an autophagosome.

### The initiation of autophagosome formation is inefficient

Previous work by others has largely focused on determining the number of cytoplasmic autophagy factor foci as a measurement of autophagosome formation, and a limited number of studies have analyzed the lifetime of foci formed by autophagy proteins (Dalle Pezze et al., 2021; Karanasios et al., 2013a; Karanasios et al., 2013b). Our dual color imaging of the tagged autophagy proteins and P62, which marks phagophores that are in the process of cargo sequestration and mature autophagosomes, allowed us to systematically analyze the kinetics of phagophore maturation. Surprisingly, approximately 90% of the foci formed by ULK1, ATG13, WIPI2, ATG5, and ATG16 do not proceed to accumulate detectable quantities of P62. In addition, autophagy protein foci that do not co-localize with P62 are very short-lived (mean = 25 seconds), compared to foci that proceed to accumulate P62 (mean = 125 seconds). These observations suggest that these short-lived foci represent aborted autophagosomes that initiate the accumulation of the ULK1 complex, WIPI2, and the LC3 lipidation machinery, but rapidly disassembly rather than maturing into an autophagosome. It is important to note that our observation that 90% of ULK1, ATG13, WIPI2, ATG5, and ATG16 foci do not proceed to accumulate P62 is likely an overestimation. Our methodology to track P62 and autophagy factor foci and to determine their co-localization is very stringent and we likely fail to detect all co-localized trajectories. Strikingly, our analysis of the maturation of ATG2 foci revealed that they are twice as likely to co-localize with P62 compared to the other autophagy factors analyzed (Fig. 6D). As discussed above, the number of ATG2 foci formed is approximately half of the number formed by ULK1, ATG13, ATG5, and ATG16. Therefore, the total number of foci formed by all autophagy factors analyzed that proceed to co-localize with P62 is comparable. Together these observations suggest that the recruitment of ATG2 and the concomitant tethering to donor membranes is a critical step in committing ATG9 vesicles towards maturation into an autophagosome (Fig. 7F).

### Overall maturation kinetics of autophagosomes in human cancer cells

Foci formed by autophagy proteins have been extensively used to infer the lifetime of autophagosomes (Dalle Pezze et al., 2021; Fujita et al., 2008; Karanasios et al., 2013a; Karanasios et al., 2016; Stavoe et al., 2019). Current estimates of autophagosome lifetime rely upon manual identification and analysis of foci formed by stably expressed, fluorescently tagged autophagy protein transgenes (Dalle Pezze et al., 2021; Karanasios et al., 2013a). Using our cell lines that express fluorescently tagged autophagy proteins from their endogenous loci avoids known artifacts due to protein overexpression and provides comparable experimental conditions to directly compare individual autophagy factors. In addition, our observations described above demonstrate that a large fraction of autophagy protein foci does not mature into autophagosomes. To accurately analyze the kinetics of autophagosome formation it is, therefore, necessary to combine the detection of autophagy factor foci with a marker of mature autophagosomes, such as P62 or LC3B. In addition, our fully automated detection and particle tracking approach provide an unbiased method that generates a tremendous amount of data compared to previous methodologies that relied on manual tracking of a small number of foci (Dalle Pezze et al., 2021; Karanasios et al., 2013b).

Our observations demonstrated that all autophagy factors analyzed accumulate approximately 40 seconds prior to P62 recruitment. This suggests that ULK1, ATG13, ATG5, ATG16, WIPI2, and ATG2 are all recruited in rapid succession (Fig. 7F). Our current imaging approach does not have the time resolution required to determine a precise recruitment order of the autophagy factors analyzed. In general, we note that previous analysis of foci report longer lasting values than our data shows. For instance, previous analysis of ATG5 and ATG13 recruitment kinetics showed ATG5 foci lasting 480 seconds and ATG13 foci which lasted for 200 seconds on average, compared to the average lifetime of 102 and 82 seconds respectively that we observed in our experiments (Dalle Pezze et al., 2021; Karanasios et al., 2013a; Koyama-Honda et al., 2013). The potential discrepancy may be due to manual notation, overexpression effects, or cell line differences.

Once P62 accumulation is detectable the autophagy factors remain associated with the phagophore for 50-80 seconds, and P62 signal is detected for 60-90 seconds after the autophagy proteins have dissociated from the autophagosome. As an additional control, we used LC3B instead of P62 as a marker for mature autophagosomes which lead to similar results. Overall, similar to ATG protein foci length, the lifetime of LC3/P62 positive autophagosomes are substantially shorter than previously reported in the literature (Dalle Pezze et al., 2021; Karanasios et al., 2013a; Karanasios et al., 2013b). We assume that the disappearance of the P62 signal is a consequence of the fusion of the autophagosome with the lysosome, which quenches the fluorescent signal of GFP. In total, these experiments demonstrate that the average lifetime of autophagosome biogensesis from its initiation is 110 seconds (ranging from 90-120 seconds) in U2OS cells. Importantly, the autophagosome lifetime we determined using automated tracking of thousands of autophagy foci was comparable in all HaloTagged autophagy protein cell lines, raising our confidence that we are measuring the lifetime time of bona fide autophagosomes. The similarity in the overall lifetime also indicates that the autophagosome formation kinetics are not adversely affected by the HaloTag fusion proteins. Finally, our observations demonstrate that all autophagy factors analyzed are removed from autophagosomes prior to the fusion of the autophagosome with the lysosome. This suggests that the signal that leads to the recruitment of these proteins, likely the enrichment of PI3P, is removed from the autophagosome once it has closed and is fully matured (Fig. 7F).

In total, these experiments precisely define the overall lifetime of autophagosomes, the timing of autophagosome maturation, and the time it takes for a mature autophagosome to fuse with a lysosome. Importantly, we frequently observed P62 foci that displayed rapid movements along linear trajectories, especially after P62 signal had been present for an extended period of time. This suggests that autophagosomes or maturing phagophores are transported by cytoskeletal motors, potentially to capture cargo or to accelerate their collision with a lysosome.

Altogether our work significantly expands our mechanistic un derstanding of autophagosome formation and provides a sophisticated experimental framework and toolkit for future quantitative studies of autophagy in human cancer cells.

## ACKNOWLEDGEMENTS

The order of authors D.B. and C.B. was decided by a randomization process and both authors contributed equally to the paper; co-first authors reserve the right to list themselves first on their curriculum vitae. This work was supported by a grant from the NIH (DP2 GM142307) to J.C.S. J.C.S. was a Damon Runyon Dale F. Frey Scientist supported (in part) by the Damon Runyon Cancer Research Foundation (DFS-24-17). We thank Dr. Daniel T. Youmans and Dr. Thomas R. Cech for providing the plasmid for recombinant production of the 3xFLAG-HaloTag protein. We thank Dr. Eric Patrick for contributing to the preparation of the recombinant HaloTag protein used in this study. Fig.1A, Fig.S1A, and Fig.S4A were created in or adapted from BioRender. This preprint was generated using a [template](https://www.overleaf.com/latex/templates/henriqueslab-biorxiv-template/nyprsybwffws) from Ricardo Henriques.

## AUTHOR CONTRIBUTIONS

Conceptualization: D.B., C.B., and J.C.S.; Experiments: D.B. and C.B.; Data Analysis: D.B. and C.B.; Writing – Original Draft: D.B. and C.B.; Writing – Review and Editing: D.B., C.B., and J.C.S.

## EXPERIMENTAL MODEL AND SUBJECT DETAILS

### Cell Lines and Tissue Culture

The cell lines were derived from human bone osteosarcoma epithelial cells (U2OS line) and grown in RPMI cell culture media supplemented with 10% fetal bovine serum (FBS), 100 units/ml penicillin, 100 µg/ml streptomycin at 37°C with 5% CO_2_.

## METHOD DETAILS

### Plasmid Construction and Genome Editing

The autophagy genes were edited at their endogenous 5’-end, except for ULK1, which was edited at the 3’ end. For the 5’-end editing, the 3XFLAG-HaloTag donor plasmid was generated according to the procedure described by Xi et al. (Schmidt et al., 2016; Xi et al., 2015). Between the HaloTag and the exon-1 sequence, a linker sequence including a TEV protease cleavage site was inserted. The homology arms and HaloTag fragments were ligated into pFASTBac linearized with HpaI using Gibson Assembly (NEB). All single-guide RNAs (sgRNAs) were cloned into pX330 Cas9 donor plasmid as previously described (Cong et al., 2013). Cells were transfected using the FuGENE HD® transfection reagent (Promega Co.); after 72h of transfection, the cells were selected for 4-6 days with puromycin (1µg/ml). Following selection with puromycin, cells were transfected with an eGFP-CRE recombinase plasmid. Cells were then labeled with the HaloTag ligand JF646 (Janelia) and sorted with FACS using the eGFP/JF646 positive signals. In single-cell clones, homologous recombination was confirmed by PCR and Sanger sequencing. All PCR oligonucleotides and sgRNA sequences are listed in the Resource Table.

### Western Blotting

Gene-editing protein expression compared to parental U2OS was assessed by western blotting. The protein samples were separated on 4–15% Bis-Tris gels (Life Technologies), followed by standard western blotting procedures. The primary and secondary antibodies used are listed in the Resource Table. Clarity western ELC Substrate (BioRad) was used to generate a chemiluminescence signal detected with a ChemiDoc imaging system (BioRad). Expression levels of the edited cell lines were estimated by comparing the chemiluminescence signal with the parental U2OS after normalization with the stain-free signal.

### LC3 assay

Cells were grown in 24-well plates at 60-70% confluency and treated with 100nM rapamycin and 100nM rapamycin + 100nM bafilomycin for 2h; a control in regular RPMI media was also performed. After treatment, the cells were harvested with 150µl sample buffer, and 20ul were loaded in a 4-20% Bis-Tris gel. Gels were transferred into PVDF blot and treated against LC3 antibody according to the protocol supplied by the provider (CellSignaling). Chemiluminescence signal was detected with a ChemiDoc imaging system (BioRad) and signal corresponding to LC3-I andLC3-II was quantified using the ImageQuant™ software (Cytiva).

### Autophagy proteins and LC3 foci quantification

Cells (10,000) were grown in 96-well optical plates and transfected after 24h with GFP-LC3 using the viral BacMam 2.0 transfection reagent at a concentration of 0.25 × 108 viral particles/mL. After 24h, cells were labeled with 100nM JF646 for 10 min, followed by 3 washed with growth media and 5min bleeding at 37°C. Nuclear stain (Hoechst dye, diluted 1:10000) was added during the last wash step. For starvation, cells were washed 3 times with PBS buffer before the addition of EBSS starvation media. Cells were imaged on an Olympus microscope equipped with a X-Cite TURBO LaserLED illumination system (Excelitas Technologies), on a 60X objective. For most of the cell lines, the lasers were set at the following intensity power: 650nm, 100% intensity; 488nm, 30% intensity; 385nm, 5% intensity. For avoiding signal saturation, for Halo-LC3 the 640nm lased was set at 20% intensity power. For each cell line, a single frame was taken at 100ms exposure for the three channels. Images were processed using a home-built foci analysis algorithm written in Icy (BioImage Analysis Lab).

### Expression and Purification of Recombinant His-3XFlag-HaloTag

The His-3X-FlagHaloTag construct kindly provided by Dr. Youmans and Dr. Cech at the University of Colorado. HaloTag protein was expressed and purified from E. coli BL21(DE3) cells grown in LB medium. Cells were grown at 37°C and expression was induced at OD~0.5-0.8 with 1mL of IPTG (1M). Upon induction, the temperature was decreased at 18°C, and cells were grown overnight (16h) before harvesting. Cells were centrifugated, the pellet resuspended in wash buffer (50 mM sodium phosphate buffer pH 8.0, 300 mM sodium chloride, 10 mM imidazole, 5 mM beta-mercaptoethanol), and sonicated for 2 min (40% amplitude, 90 seconds of sonication, 10 second pulses, 20 second pause, FisherbrandTM Model 505, 0.5 inch tip). The lysate was then centrifuged (40,000g and 4°C for 30min) and the supernatant loaded to 1 mL of in fast flow nickel Sepharose (Cytiva) and incubated under gentle rotation for 1h at 4°C. Beads were then washed 3 times with 6ml of wash buffer, and elution was performed using elution buffer (0 mM sodium phosphate buffer pH 7.0, 300 mM sodium chloride, 250 mM imidazole, 5 mM beta-mercaptoethanol). The eluate was subjected to further purification using size-exclusion chromatography on a Superdex S75 column in isocratic mode (50 mM Tris pH 7.5, 150 mM potassium chloride, 1 mM DTT). Protein was concentrated to 1mg/ml, supplemented with 50% glycerol, flash-frozen, and stored at -80°C. To fluorescently label the 6xHIS-3xFLAG-HaloTag, the protein was incubated with a 2-fold excess of JF646 HaloTag-ligand overnight at room temperature. Excess fluorescent dye was removed by size exclusion chromatography using a Superdex S75 column in isocratic mode (50 mM Tris pH 7.5, 150 mM potassium chloride, 1 mM DTT). The protein concentration and labeling efficiency were determined by absorption spectroscopy using λ_280nm_ = 41,060 M^-1^ cm^-1^ for the 6xHIS-3xFLAG-HaloTag and λ_646nm_ = 152,000 M^-1^ cm^-1^ for the JF646 fluorescent dye (Grimm et al., 2015).

### In-Gel Fluorescence Absolute Protein Quantification

His-3XFlag-HaloTag was labeled by combining 15 nmol of JF646 (30ul from 0.5mM DMSO resuspended stock) to 0.2 mg recombinant protein in 200µl of buffer and incubated at RT for 30 minutes. The labeled protein was then purified by size exclusion on a S75 column. Fractions were combined and concentrated in Vivaspin 10 kDa columns and glycerol was added for a final concentration of 20%. Vials were snapfrozen in liquid nitrogen and stored at -80°C. The final concentration was measured using a 40ul microcuvette in a fluorescence spectrophotometer. Protein concentration was calculated using an extinction coefficient ε =152 mM^-1^ cm^-1^ (Grimm et al., 2017). The labeled tag was then serial diluted in sample buffer from 37.5 fmol down to 0.29 fmol and added to cell lysates that were prepared using direct lysis in sample buffer and diluted from 150,000 cells/mL to 75,000 cells/mL in 25,000 cell/mL increments. The standards were then aliquoted into individual use tubes, boiled, snap-frozen in liquid nitrogen, and stored at -80°C. For the in-gel fluorescence, 300,000 cells were seeded 24h before. The protein and the standard were loaded and separated on 4–15% BisTris gels. Fluorescence was detected using the Cy5.5 filter on a Sapphire™ molecular imager (Azure Biosystems Inc.). The gels were quantified using the ImageQuant™ software (Cytiva).

### Flow cytometry protein abundance

The day before, 80,000 cells were plated into a 24-well plate. Cells were then labeled with JF646 at 500nM for 30 minutes in media and washed with a 5-minute incubation in media. Cells were then harvested and transferred to a 2 ml deep 96-well plate using a multichannel pipette and 5mM EDTA in PBS. PFA fixation solution was added for a final concentration of 2% and incubated for 10 minutes. Samples were then washed in 1% BSA in PBS once and filtered into a 96-well round-bottom plate. Analysis was performed on a Cytek Aurora using a 96-well loader and unmixed using the default SpectroFlo software.

### Single-Molecule Live-Cell Imaging

We performed singleparticle tracking analysis for determining diffusion coefficients of HaloTagged autophagy protein under control and EBSS starving conditions. Single-molecule live-cell imaging was carried out on an Olympus microscope, equipped with a TIRF/HiLo illumination system and four laser lines (405nm, 488nm, 561nm, and 647nm). The microscope is equipped with an environmental chamber (cellVivo) to control humidity, temperature, and CO2 level, two iXon Ultra 897 EMCCD cameras (Andor), a 100x TIRF oil-immersion objective (Olympus UApo), and appropriate excitation and emission filters. Cells (200-300k) were grown on glass coverslips (170± 5 µm, Schott) and imaged after ~24h. Coverslips were cleaned with 1M KOH (1h in a sonicated water bath), rinsed with ddH2O, cleaned with 100% EtOH (1h in a sonicated water bath), and dried under N2 stream before assembly on 35mm diameter imaging dish. Precise determination of diffusion coefficient using single-particle tracking requires sparse labeling of the Halo-tagged protein. Cells were labeled with Halo ligand JFX650 with the following optimized conditions: 1.0 nM, 30s pulse (ATG9); 500pM, 30s pulse (WIPI2, ULK1, ATG2, ATG5, ATG13, ATG16L1, PI4KIIIβ); 50pM, 30s pulse (LC3). These labeling concentrations allowed a maximum of 10-15 particles per frame through the entire imaging experiment (3000 frames, 20.4s per movie). After labeling, the cells were washed three times with fresh media and incubated for an additional 10 min at 37°C (5% CO2). To avoid additional labeling due to residual Halo ligand, we block non-labeled HaloTag with 7-bromophenol (10µM) in both control media and EBSS. Before adding EBSS, cells were washed 3X with PBS. EBSS-treated samples were imaged within one hour of treatment. To image autophagy proteins, cells were imaged continuously with the 647 nm laser (~25% laser power), with 6.8 ms exposure time, achieving 146 frames per s, using a 140×512 pixel region of interest.

### Diffusion coefficient determination

For detecting individual single-molecule particles and compute particle tracks, we used the batch parallel-processing version of SLIMFast in MatLab 2020b. This version – kindly provided by Xavier Darzacq and Anders Hansen – allows direct import of the TIFF files and uses the MTT algorithm (Sergé et al., 2008). For tracking, the following settings were used for all the proteins: Exposure Time = 6.8 ms, NA = 1.49, PixelSize = 0.16 µm, Emission Wavelength = 664 nm, Dmax= 15 µm^2^ s-1, Number of gaps allowed = 2, Localization Error = 10-5, Deflation loops = 0. For ATG9, the Dmax was set at 5 µm^2^ s-1. From the tracked particles, we determined the diffusion coefficients and the fraction corresponding to distinct particle subpopulation using the MATLAB version of SpotOn (kindly provided by Xavier Darzacq and Anders Hansen) (Hansen et al., 2018). The following settings were used: TimeGap = 6.8 ms, dZ = 0.700 µm, GapsAllowed = 2, TimePoints = 7, JumpsToConsider = 4, BinWidth = 0.01 µm, PDFfitting, D_Free1_3State = [2 15], D_Free2_3State = [0.15 2], D_Bound_3State = [0.0001 0.15]. For ATG9, a 2-state model was applied, with the following settings: D_Free2_1State = [0.2 5], D_Bound2_State = [0.0001 0.2]. We carried out 3 independent biological replicates with at least 20 cells for each cell line for all experiments.

### Determination of foci lifetime and intensity for HaloTag autophagy proteins

Foci lifetimes were determined in the HaloTag edited by live-cell imaging. As for the singlemolecule imaging, foci determination was carried out on the Olympus microscope, using a 60X oil immersion objective (Olympus UApo), and appropriate excitation and emission filters. The illumination was achieved with a X-Cite TURBO LaserLED illumination system (Excelitas Technologies), equipped with a 650 LED source set a 100% intensity power. Cells (150k) were grown on glass coverslips (170 ± 5 µm, Schott) and imaged after 48h. Cells were labeled with Halo ligand 100nM JF646 for 10min, which allows quantitative labeling of HaloTagged proteins. After labeling, the cells were washed three times with fresh media and incubated for an additional 10 min at 37°C (5% CO_2_). To avoid additional labeling due to residual Halo ligand, we blocked non-labeled HaloTag with 7-bromophenol (10µM) in both control media and EBSS. Before adding EBSS, cells were washed 3X with PBS. EBSS-treated samples were imaged within one hour of treatment. For each condition, four cell clusters (4-6 cells x cluster) were selected and imaged at 15s-time intervals (50ms exposure time) over 1 hour, corresponding to 240 frames per movie. For ATG2 and ATG13, cells were also imaged at a faster rate (3s) for over 240 frames (corresponding to 12 min total imaging). The experiments were run in triplicate, with approximately 20-30 cells imaged per cell line. Foci were treated as single-particles and analyzed using TrackIT with the following settings: threshold 1.5; tracking radius 6; minimum track length 2; gap frames 2. For calculating foci intensity, we centered a 9×9 pxl box on the particle coordinated obtained by TrackIt. We then divided this box in two subregions. The central 5×5 pxl box – corresponding to the foci signal – was used to obtain the intensity of the particle as the max intensity (I_max_). The pixel intensity of the remaining subregion surrounding the 5×5 pxl area was averaged to compute the background intensity (I_background_). The corrected particle intensity (I_max, corr_) was calculated as following I_max, corr_ = I_max_ I_background_. Finally, for each foci track, we calculated the foci intensity by averaging the I_max, corr_ associated with each particle.

### Determination of foci colocalization with P62 and LC3 foci

Cells were seeded at low confluency (100k) on glass coverslips (170 ± 5 µm, Schott). After 24h, cells were transduced with GFP-LC3 or GFP-P62 using the viral BacMam 2.0 transfection reagent at a concentration of .25 × 108 viral particles/mL. After an additional 24h from viral transduction, cells were labeled for foci visualization (see the previous section) and imaged in control and EBSS starvation media. Before adding EBSS, cells were washed 3X with PBS. EBSS-treated samples were imaged within one hour of treatment. Cells were imaged on the Olympus microscope, using a 60X oil immersion objective (Olympus UApo), and appropriate excitation and emission filters. The illumination was achieved with an X-Cite TURBO LaserLED illumination system (Excelitas Technologies), equipped with a 650 nm (100% intensity power) and a 488 nm (30% intensity power) LED source. For each condition, four cell clusters (4-6 cells x cluster) were selected and imaged at 3s time interval (50ms exposure time) over 12 min, corresponding to 240 frames per movie. The experiments were run in triplicate, with approximately 20-30 cells imaged per cell line. Autophagy protein foci were treated as single particles and analyzed using TrackIT with the following settings: threshold 1.5; tracking radius 6; minimum track length 2; gap frames 6. Tracks were exported and analyzed with a custom-built Matlab (v. 2022a) algorithm. The algorithm calculated the Euclidean distance between each tracked autophagy foci with the nearest P62 (or LC3) foci. If the calculated distance was 3 pxl (0.81 µm), the P62 particle was considered colocalized with the tracked autophagy protein foci. An ATGs foci track was considered colocalized with a P62 (or LC3) foci track if at least 4 colocalization events were recorded. The colocalization between ATGs and P62 (or LC3) foci was computed by calculating the difference between the first and last colocalization events. Diffusion coefficients for the autophagy foci were computed using MSDAnalyzer implemented in MatLab or built in the TrackIt package. In both cases, the algorithm fit the mean-squared displacement (MSD) curve obtained from the SPT with a power law equation describing anomalous diffusion. For diffusive analysis, only 60% of the track was fitted.

### STATISTICAL ANALYSIS

#### Quantification of Western Blots

Gel images from Western Blots were analyzed using ImageQuant TL 8.2. The statistical significance of the observed differences was calculated using a two-tailed T-test using a minimum of three biological replicates. Each biological replicate was analyzed in technical triplicate.

#### Statistical Analysis

Data are expressed as ± one standard deviation (SD) unless otherwise stated. Statistical tests were performed as stated.

## Supplementary Movie Legends

**Movie S1**. Representative live-cell single-molecule imaging movies of control and EBSS-treated U2OS cells expressing 3xFLAG-HaloTagged ATG2A labeled with JFX650 and acquired at 146 frames per second. 140×140 pixels with a pixel size of 0.16 µm.

**Movie S2**. Representative live-cell single-molecule imaging movies of control and EBSS-treated U2OS cells expressing 3xFLAG-HaloTagged ATG5 labeled with JFX650 and acquired at 146 frames per second. 140×140 pixels with a pixel size of 0.16 µm.

**Movie S3**. Representative live-cell single-molecule imaging movies of control and EBSS-treated U2OS cells expressing 3xFLAG-HaloTagged ATG9A labeled with JFX650 and acquired at 146 frames per second. 140×140 pixels with a pixel size of 0.16 µm.

**Movie S4**. Representative live-cell single-molecule imaging movies of control and EBSS-treated U2OS cells expressing 3xFLAG-HaloTagged ATG13 labeled with JFX650 and acquired at 146 frames per second. 140×140 pixels with a pixel size of 0.16 µm.

**Movie S5**. Representative live-cell single-molecule imaging movies of control and EBSS-treated U2OS cells expressing 3xFLAG-HaloTagged ATG16 labeled with JFX650 and acquired at 146 frames per second. 140×140 pixels with a pixel size of 0.16 µm.

**Movie S6**. Representative live-cell single-molecule imaging movies of control and EBSS-treated U2OS cells expressing 3xFLAG-HaloTagged ULK1 labeled with JFX650 and acquired at 146 frames per second. 140×140 pixels with a pixel size of 0.16 µm.

**Movie S7**. Representative live-cell single-molecule imaging movies of control and EBSS-treated U2OS cells expressing 3xFLAG-HaloTagged WIPI2 labeled with JFX650 and acquired at 146 frames per second. 140×140 pixels with a pixel size of 0.16 µm.

**Movie S8**. Representative live-cell single-molecule imaging movies of control and EBSS-treated U2OS cells expressing 3xFLAG-HaloTagged LC3B labeled with JFX650 and acquired at 146 frames per second. 140×140 pixels with a pixel size of 0.16 µm.

**Movie S9**. Representative live-cell single-molecule imaging movies of control and EBSS-treated U2OS cells expressing 3xFLAG-HaloTagged PI4K3β labeled with JFX650 and acquired at 146 frames per second. 140×140 pixels with a pixel size of 0.16 µm.

**Movie S10**. Representative live-cell single-molecule imaging movies of control and EBSS-treated U2OS cells expressing 3xFLAG-HaloTagged NES labeled with JFX650 and SNAPTagged SEC61B labeled with JF650 and acquired at 146 frames per second. 140×140 pixels with a pixel size of 0.16 µm.

**Movie S11**. Representative live-cell imaging movies showing the automated foci detection and tracking using TrackIt algorithm. Left: raw movie of Halo-ATG2A cell line in EBSS starvation media; Center: automated identification of foci spots; Right: compiling of tracks. Cells were labeled with JF646 and acquired at 4 frames per minute. 300×300 pixels with a pixel size of 0.27 µm.

**Movie S12**. Representative live-cell foci imaging movies of control and EBSS-treated U2OS cells expressing 3xFLAG-HaloTagged ATG2A labeled with JF646 and acquired at 4 frames per minute. 300×300 pixels with a pixel size of 0.27 µm.

**Movie S13**. Representative live-cell foci imaging movies of control and EBSS-treated U2OS cells expressing 3xFLAG-HaloTagged ATG5 labeled with JF646 and acquired at 4 frames per minute. 300×300 pixels with a pixel size of 0.27 µm.

**Movie S14**. Representative live-cell foci imaging movies of control and EBSS-treated U2OS cells expressing 3xFLAG-HaloTagged ATG9A labeled with JF646 and acquired at 4 frames per minute. 300×300 pixels with a pixel size of 0.27 µm.

**Movie S15**. Representative live-cell foci imaging movies of control and EBSS-treated U2OS cells expressing 3xFLAG-HaloTagged ATG13 labeled with JF646 and acquired at 4 frames per minute. 300×300 pixels with a pixel size of 0.27 µm.

**Movie S16**. Representative live-cell foci imaging movies of control and EBSS-treated U2OS cells expressing 3xFLAG-HaloTagged ATG16 labeled with JF646 and acquired at 4 frames per minute. 300×300 pixels with a pixel size of 0.27 µm.

**Movie S17**. Representative live-cell foci imaging movies of control and EBSS-treated U2OS cells expressing 3xFLAG-HaloTagged ULK1 labeled with JF646 and acquired at 4 frames per minute. 300×300 pixels with a pixel size of 0.27 µm.

**Movie S18**. Representative live-cell foci imaging movies of control and EBSS-treated U2OS cells expressing 3xFLAG-HaloTagged WIPI2 labeled with JF646 and acquired at 4 frames per minute. 300×300 pixels with a pixel size of 0.27 µm.

**Movie S19**. Representative live-cell foci imaging movies of EBSS-treated U2OS cells expressing 3xFLAG-HaloTagged ATG13 (top left), and ULK1 (top right), FIP200 (bottom left), and ATG101 (bot-tom right) knock outs. Cells were labeled with JF646 and acquired at 4 frames per minute. 300×300 pixels with a pixel size of 0.27 µm.

**Movie S20**. Representative live-cell foci imaging movies of control and EBSS-treated U2OS cells expressing 3xFLAG-HaloTagged ATG13 with and without wortmannin. Cells were labeled with JF646 and acquired at 4 frames per minute. 300×300 pixels with a pixel size of 0.27 µm.

**Movie S21**. Representative live-cell foci imaging movies of control and EBSS-treated U2OS cells expressing 3xFLAG-HaloTagged ATG2 with and without wortmannin. Cells were labeled with JF646 and acquired at 4 frames per minute. 300×300 pixels with a pixel size of 0.27 µm.

**Movie S22**. Representative live-cell single-molecule imaging movies of control (top) and EBSS-treated (bottom) U2OS cells expressing 3xFLAG-HaloTagged ATG13 parental and and ULK1, FIP200, and ATG101 knock out. Cells were labeled with JF650 and acquired at 146 frames per second. 140×140 pixels with a pixel size of 0.16 µm.

**Movie S23**. Representative live-cell foci imaging movies of EBSS-treated U2OS cells expressing 3xFLAG-HaloTagged ATG2A la-beled with JF646 and GFP-P62 acquired at 20 frames per minute. 300×300 pixels with a pixel size of 0.27 µm.

**Movie S24**. Representative live-cell foci imaging movies of EBSS-treated U2OS cells expressing 3xFLAG-HaloTagged ATG5 la-beled with JF646 and GFP-P62 acquired at 20 frames per minute. 300×300 pixels with a pixel size of 0.27 µm.

**Movie S25**. Representative live-cell foci imaging movies of EBSS-treated U2OS cells expressing 3xFLAG-HaloTagged ATG9A la-beled with JF646 and GFP-P62 acquired at 20 frames per minute. 300×300 pixels with a pixel size of 0.27 µm.

**Movie S26**. Representative live-cell foci imaging movies of EBSS-treated U2OS cells expressing 3xFLAG-HaloTagged ATG13 la-beled with JF646 and GFP-P62 acquired at 20 frames per minute. 300×300 pixels with a pixel size of 0.27 µm.

**Movie S27**. Representative live-cell foci imaging movies of EBSS-treated U2OS cells expressing 3xFLAG-HaloTagged ATG16 la-beled with JF646 and GFP-P62 acquired at 20 frames per minute. 300×300 pixels with a pixel size of 0.27 µm.

**Movie S28**. Representative live-cell foci imaging movies of EBSS-treated U2OS cells expressing 3xFLAG-HaloTagged ULK1 la-beled with JF646 and GFP-P62 acquired at 20 frames per minute. 300×300 pixels with a pixel size of 0.27 µm.

**Movie S29**. Representative live-cell foci imaging movies of EBSS-treated U2OS cells expressing 3xFLAG-HaloTagged WIPI2 la-beled with JF646 and GFP-P62 acquired at 20 frames per minute. 300×300 pixels with a pixel size of 0.27 µm.

**Movie S30**. Representative live-cell foci imaging movies of EBSS-treated U2OS cells expressing 3xFLAG-HaloTagged ATG2A la-beled with JF646 and GFP-LC3 acquired at 20 frames per minute. 300×300 pixels with a pixel size of 0.27 µm.

**Movie S31**. Representative live-cell foci imaging movies of EBSS-treated U2OS cells expressing 3xFLAG-HaloTagged ATG13 la-beled with JF646 and GFP-LC3 acquired at 20 frames per minute. 300×300 pixels with a pixel size of 0.27 µm.

**Movie S32**. Representative live-cell foci imaging movies of EBSS-treated U2OS cells expressing 3xFLAG-HaloTagged ULK1 la-beled with JF646 and GFP-LC3 acquired at 20 frames per minute. 300×300 pixels with a pixel size of 0.27 µm.

**Movie S33**. Representative live-cell foci imaging movies of EBSS-treated U2OS cells expressing 3xFLAG-HaloTagged ATG2 (left), 3xFLAG-HaloTagged ATG2 ATG9A knock-out (right). Cells were labeled with JF646 and acquired at 4 frames per minute. 300×300 pixels with a pixel size of 0.27 µm.

**Supplementary Figure 1.**
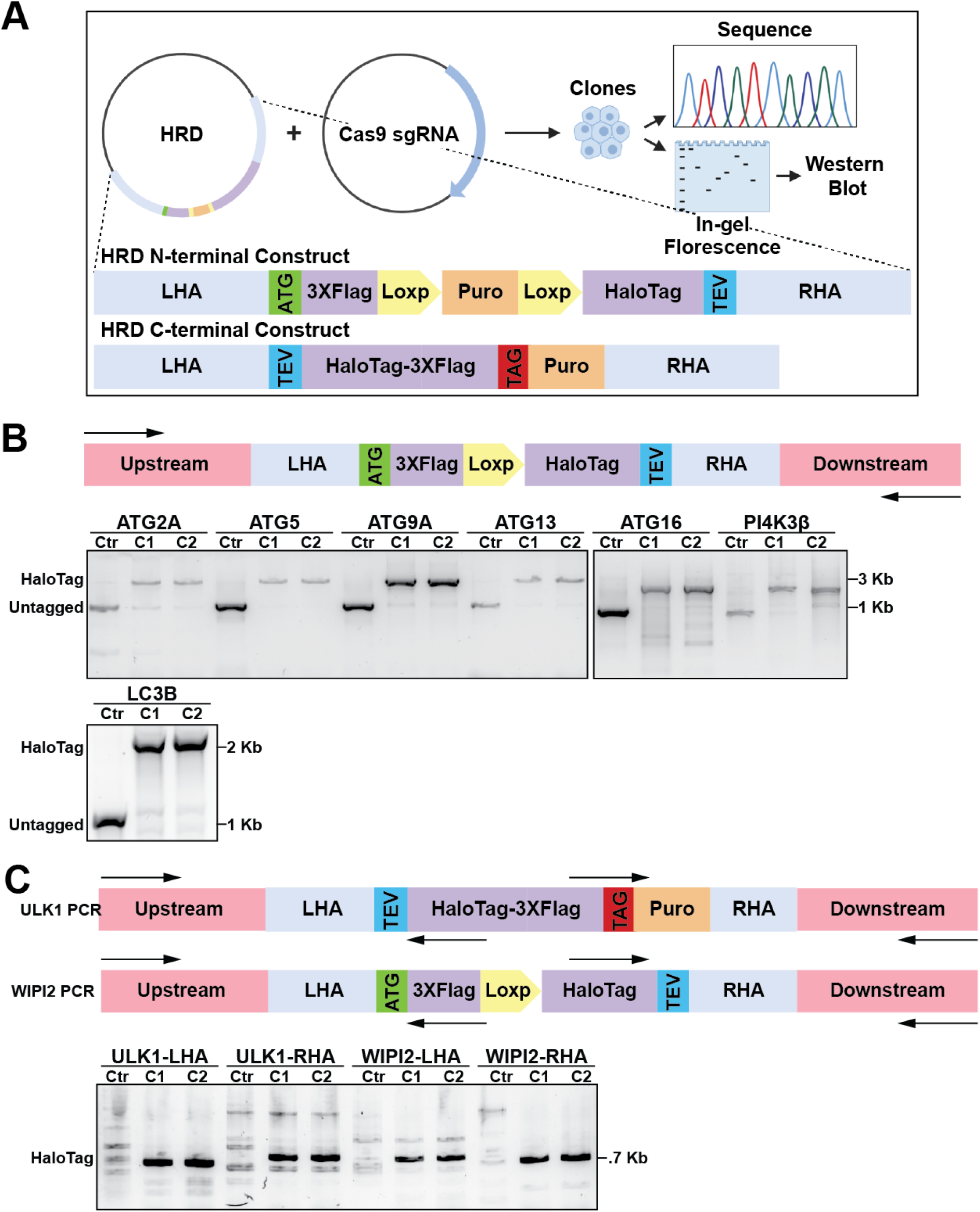
**(A)** Schematic of CRISPR-Cas9 strategy for inserting HaloTag at the N-terminus and C-terminus of autophagy genes. For N-terminus tagging, a homologous recombination donor (HDR) plasmid was supplied including the HaloTag and a puromycin resistance (Puro) cassette flanked by two LoxP sites. After puromycin selection, the Puro cassette was removed with Cre recombinase. For the C-terminus, the Puro cassette was not removed after puromycin selection. **(B)** PCR analysis from genomic DNA of genome-edited clones for verifying the correct insertion of the HaloTag at the autophagy loci. For amplifying the insertion, primers outside the homology arms region were designed. The edited clones show an expected shift of *∼*2kb on the PCR product, corresponding to the 3xFlag-HaloTag insert. (C) PCR analysis from genomic DNA of genome-edited clones for verifying the correct insertion of the HaloTag in the high GC-rich ULK1 and WIPI2 gene loci. For amplifying the insertion, primers outside the homology arms and inside the 3xFlag-HaloTag regions were designed. The edited clones show a PCR product, which is absent in the parental U2OS cell line.

**Supplementary Figure 2.**
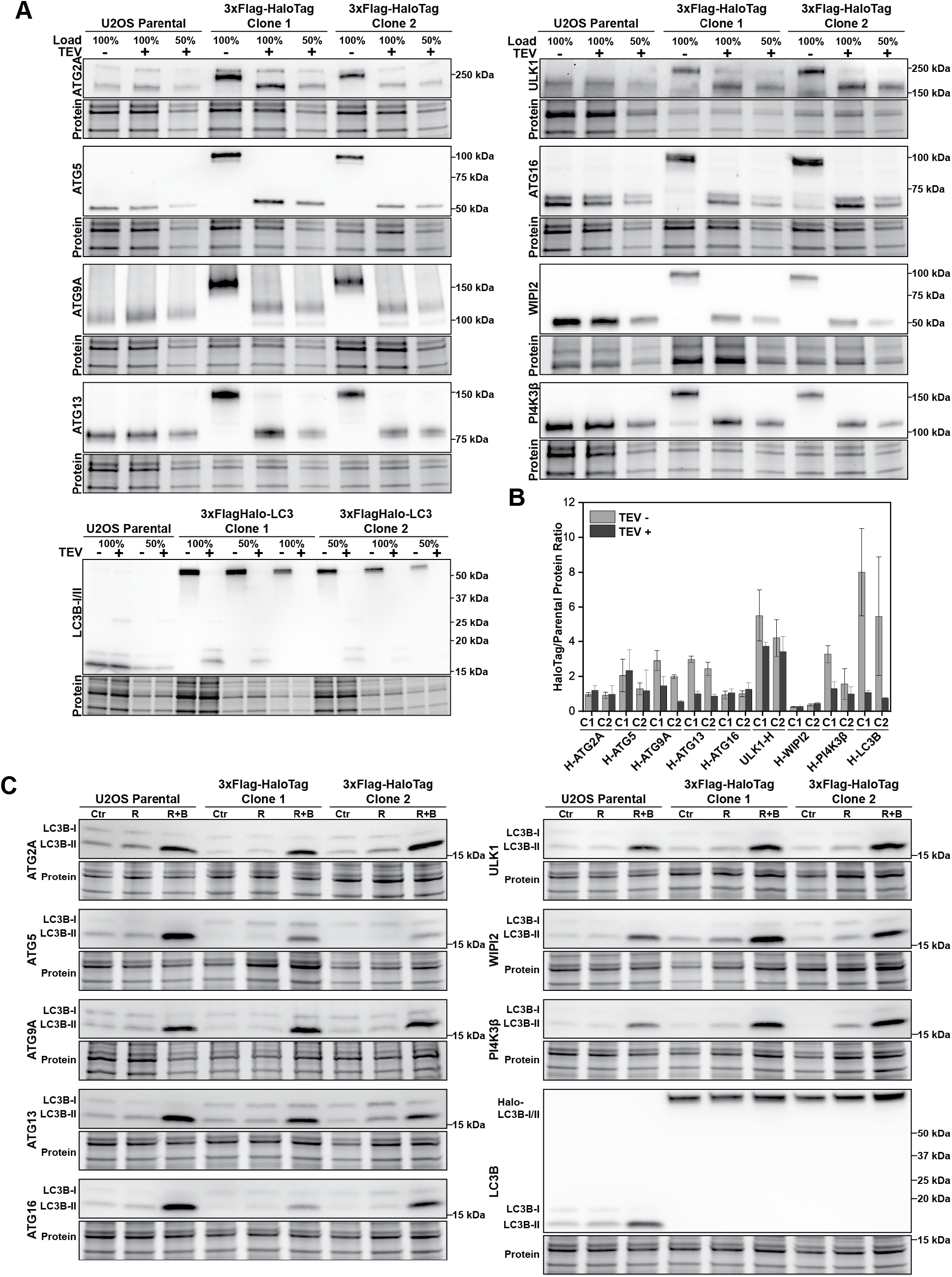
**(A)** Western blots for determining the expression levels of the HaloTagged autophagy proteins relative to the wildtype protein before and after removal of HaloTag using the TEV protease. **(B)** Quantification of the western blots (A), showing the ratio between HaloTag and parental cell line (N=3, mean ± SD). **(C)** Western blot analysis of LC3 levels in parental and genome-edited cell lines in control (Ctr) and upon treatment with rapamycin (R, 100nM for 2h), or rapamycin + bafilomycin (R+B, 100nM each, for 2h).

**Supplementary Figure 3.**
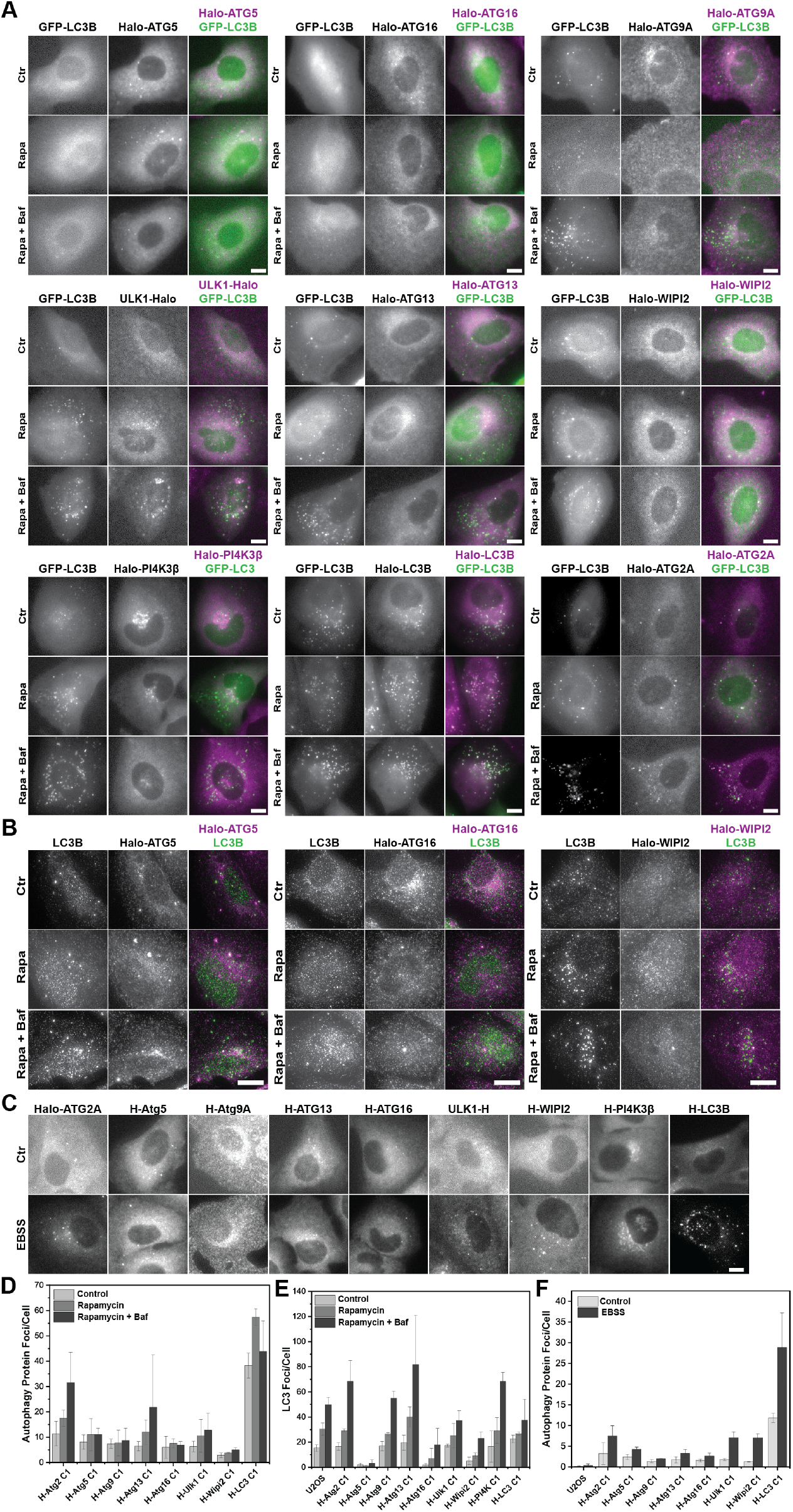
**(A)** Live-cell imaging of GFP-LC3 and JF646-labeled HaloTagged proteins in control (Ctr) and upon treatment with rapamycin (Rapa, 100nM for 2h), or rapamycin + bafilomycin (Rapa+Baf, 100nM each, for 2h). Both treatments show an expected increase in autophagy and LC3 foci. Scale bar = 10 µm. **(B)** Immunofluorescence with anti-LC3B antibody and HaloTag JF646-labeling for Halo-ATG5, Halo-ATG16, and Halo-WIPI2 cell lines in control (Ctr) and upon treatment with rapamycin (Rapa, 100nM for 2h), or rapamycin + bafilomycin (Rapa+Baf, 100nM each, for 2h). Both treatments show an expected increase in autophagy and LC3 foci. Scale bar = 10 µm. **(C)** Representative images of JF646-labeled HaloTagged proteins under control (Ctr) or EBSS starvation media, demonstrating foci-forming ability upon autophagy induction. Scale bar = 10 µm. **(D)** Quantification of autophagy foci from live-cell imaging (A). Data represent the mean ± 1SD of three biological replicates. **(E)** Quantification of LC3 foci from live-cell imaging (A) (N=3, mean ± SD). **(F)** Quantification of autophagy foci from live-cell imaging (C) (N=3, mean ± SD).

**Supplementary Figure 4.**
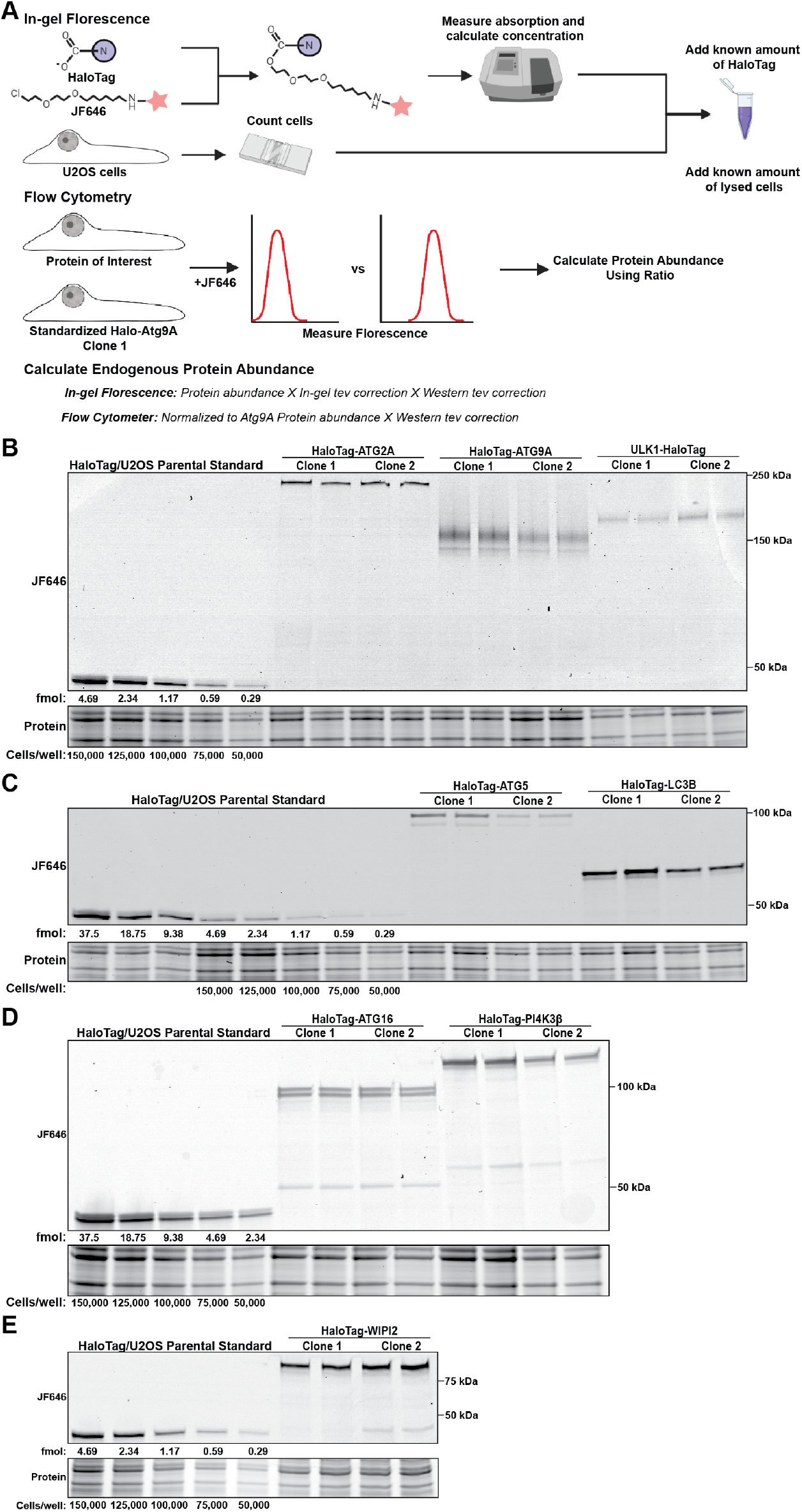
**(A)** Schematic of the protocol developed for absolute protein quantification using in-gel fluorescence (top) and flow cytometry (center). The protein copy number per cell for each method was computed using the presented equations (bottom). **(B-E)** Representative fluorescence gels for the absolute quantification of the autophagy proteins.

**Supplementary Figure 5.**
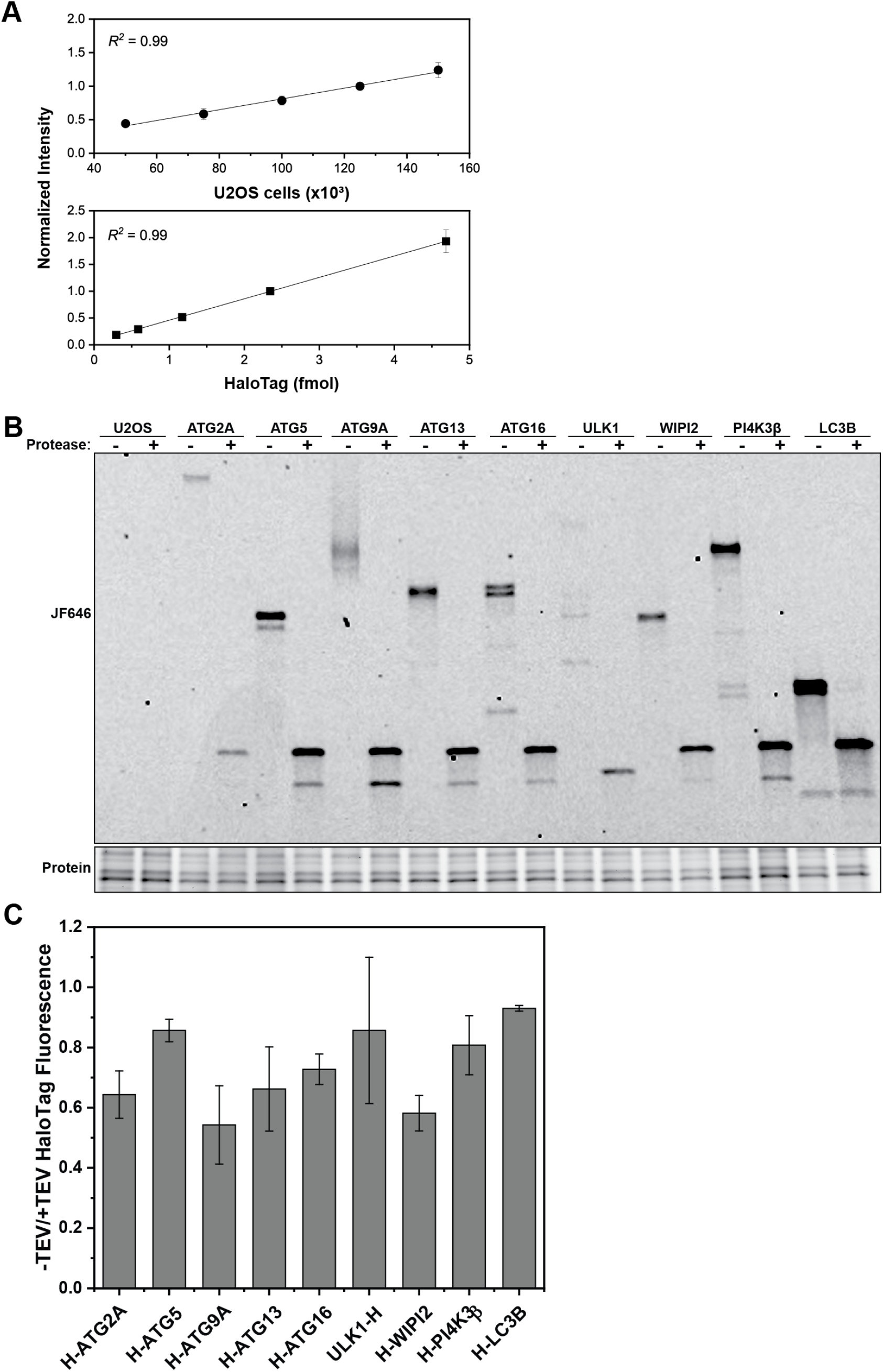
**(A)** Standard curve for cell number using stain-free gels(top) and florescent HaloTag protein (bottom), demonstrating an excellent correlation between intensity and gel loadings. **(B)** Representative fluorescence gel of HaloTagged proteins in the absence or presence of TEV protease. **(C)** Ratio of HaloTag fluorescence in the absence and presence of TEV protease (B) (N=3, mean ± SD).

**Supplementary Figure 6.**
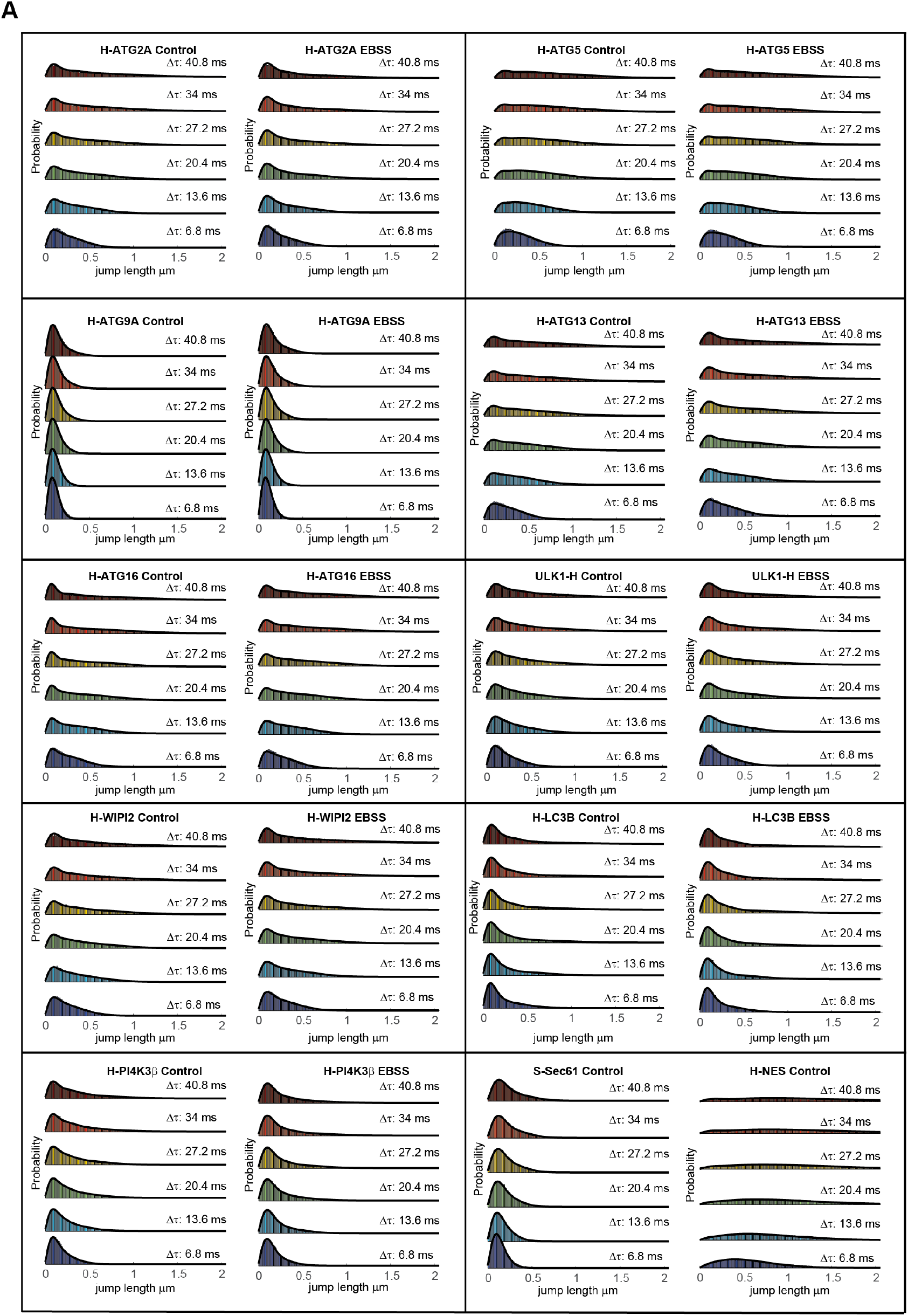
**(A)** Step-size distribution and fitting using SpotON for the HaloTagged cell lines with and without starvation. Sec61 and NES controls were run in control media only.

**Supplementary Figure 7.**
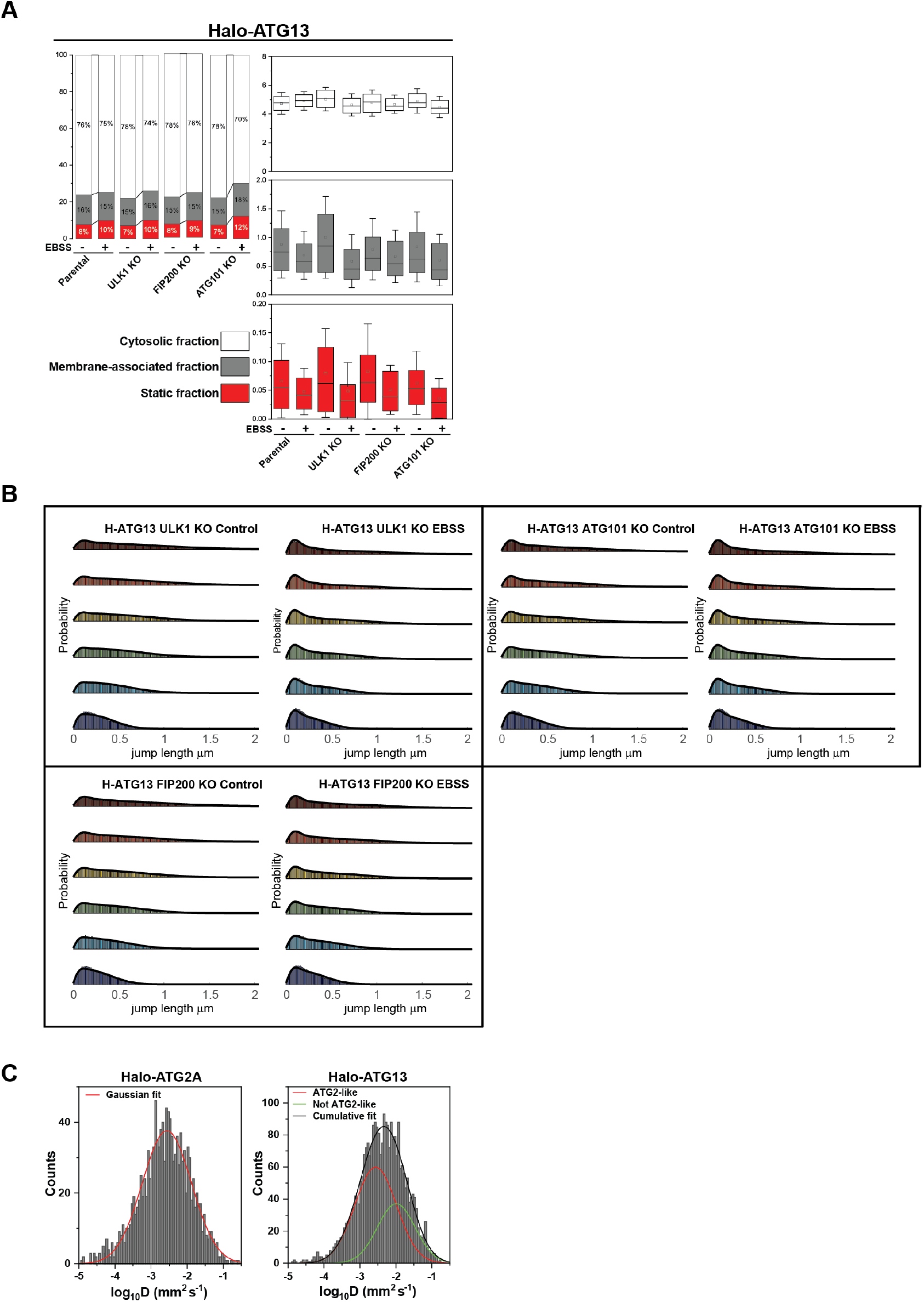
**(A)** Results of diffusive analysis for the parental Halo-ATG13 and ULK1, FIP200 and ATG101 knockout under control, and EBSS starvation. Left panel depicts the percentage associated with each fraction. Right panels present the diffusion coefficients of the tracks based on the SpotON analysis. Boxes indicate confidence interval ± SD, the square indicates the average, and the horizontal line is the median; for each condition, 3 biological replicates were analyzed, *∼*20 cells/replicate. **(B)** Step-size distribution and fitting using SpotON for parental Halo-ATG13 and knock-outs. **(C)** Distribution of diffusion coefficients for Halo-ATG2 and Halo-ATG2 foci imaged a 3s frame interval. In Halo-ATG13, Gaussian fitting shows two populations with distinct diffusive properties (solid green and red lines). Cumulative fitting is shown in a solid black line.

**Supplementary Figure.**
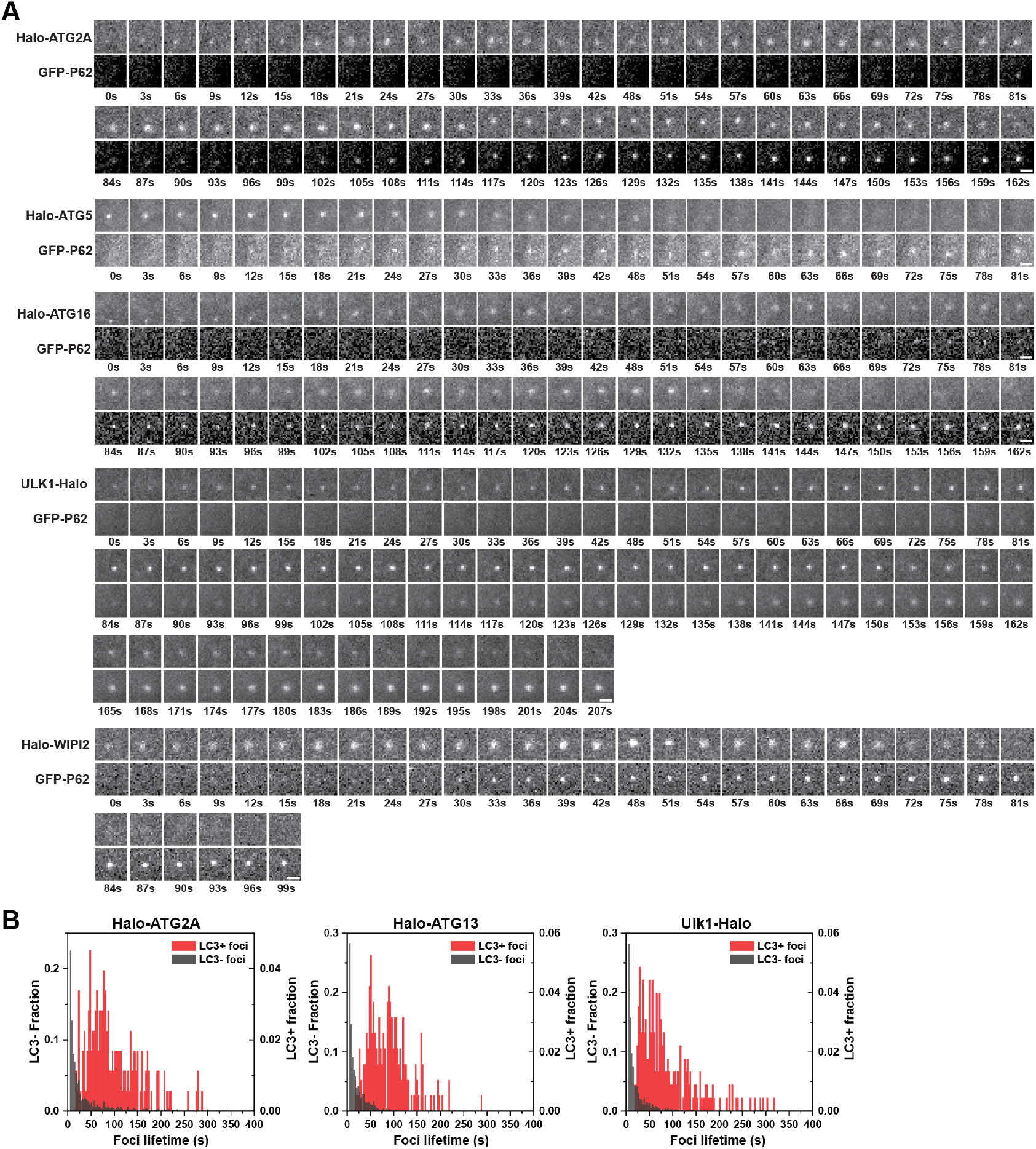
**(A)** Representative micrographs of GFP-P62 and autophagy foci. Scale bar = 10 µm. **(B)** Frequency histogram of LC3 positive (red) and LC3 negative (dark grey) autophagy foci for Halo-ATG2A, Halo-ATG13, and ULK1-Halo.

## KEY RESOURCES TABLE

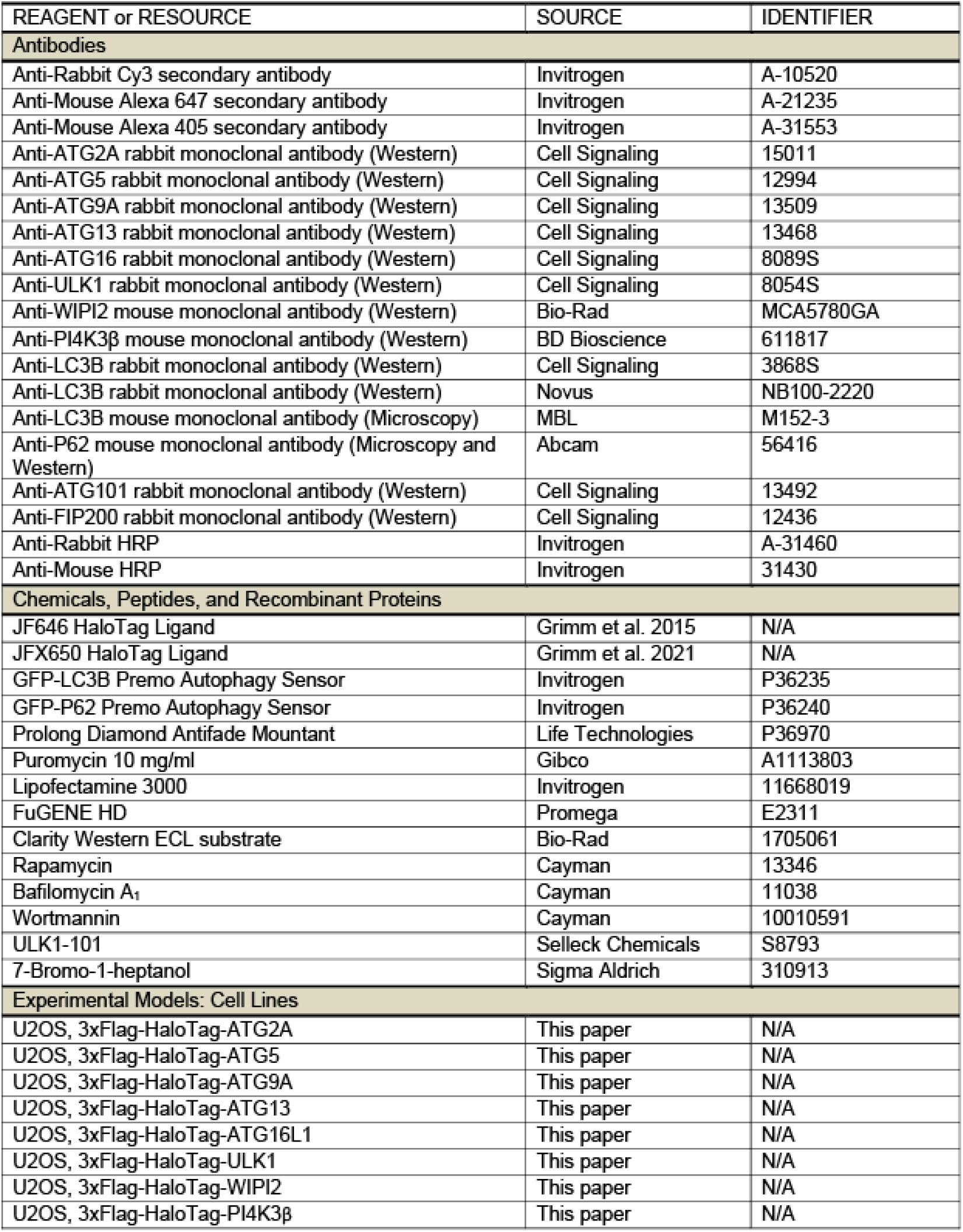
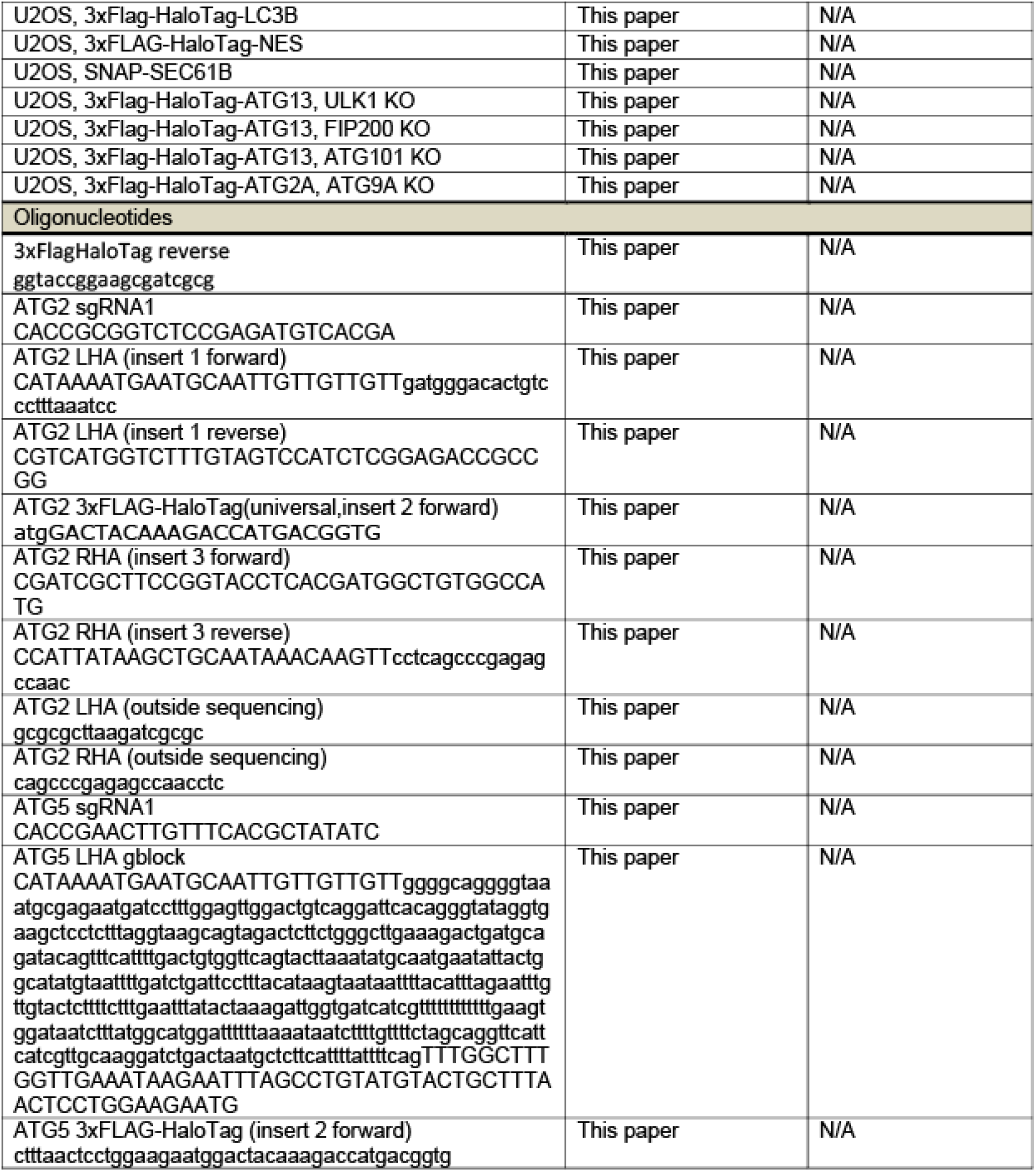
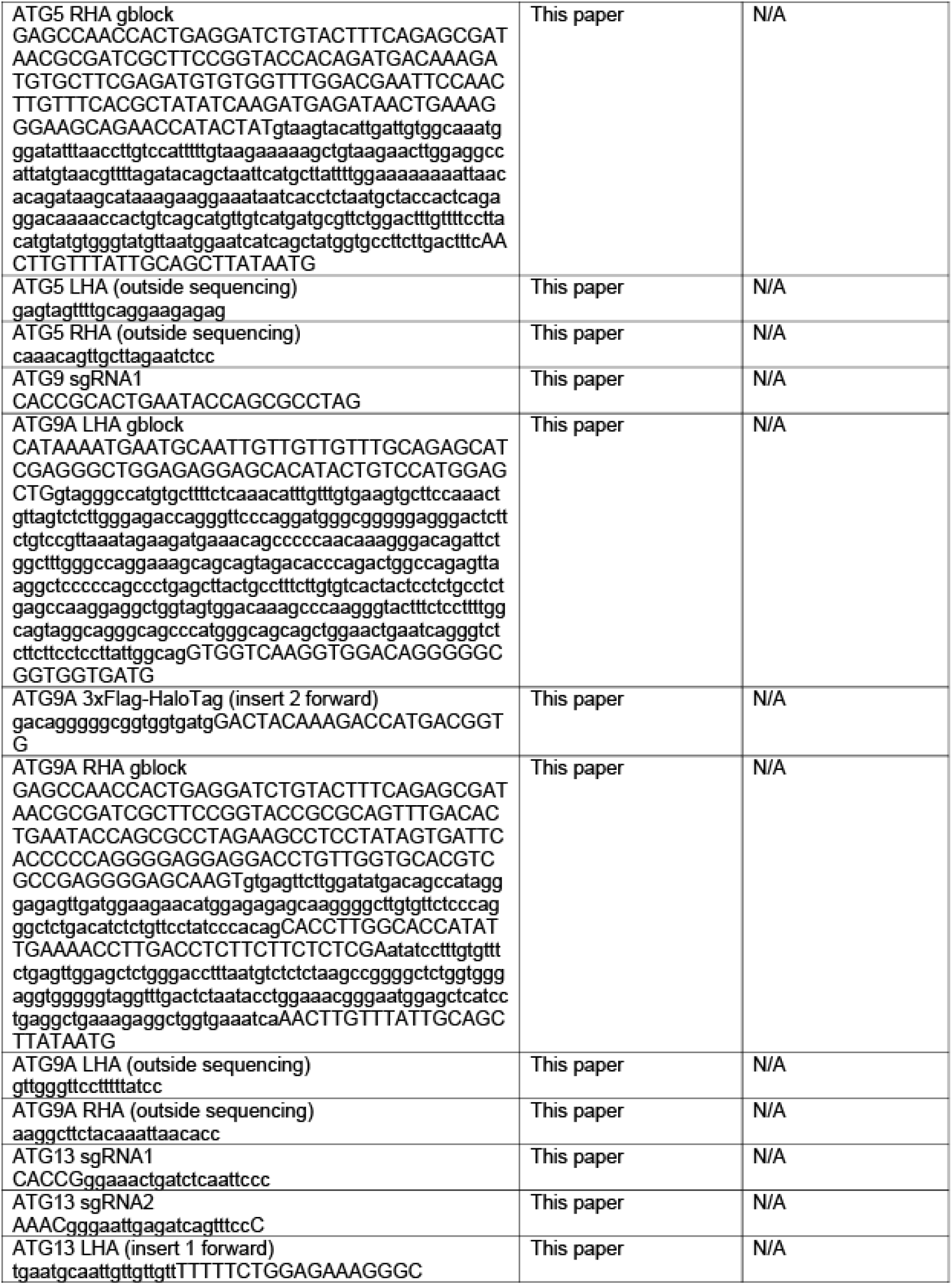
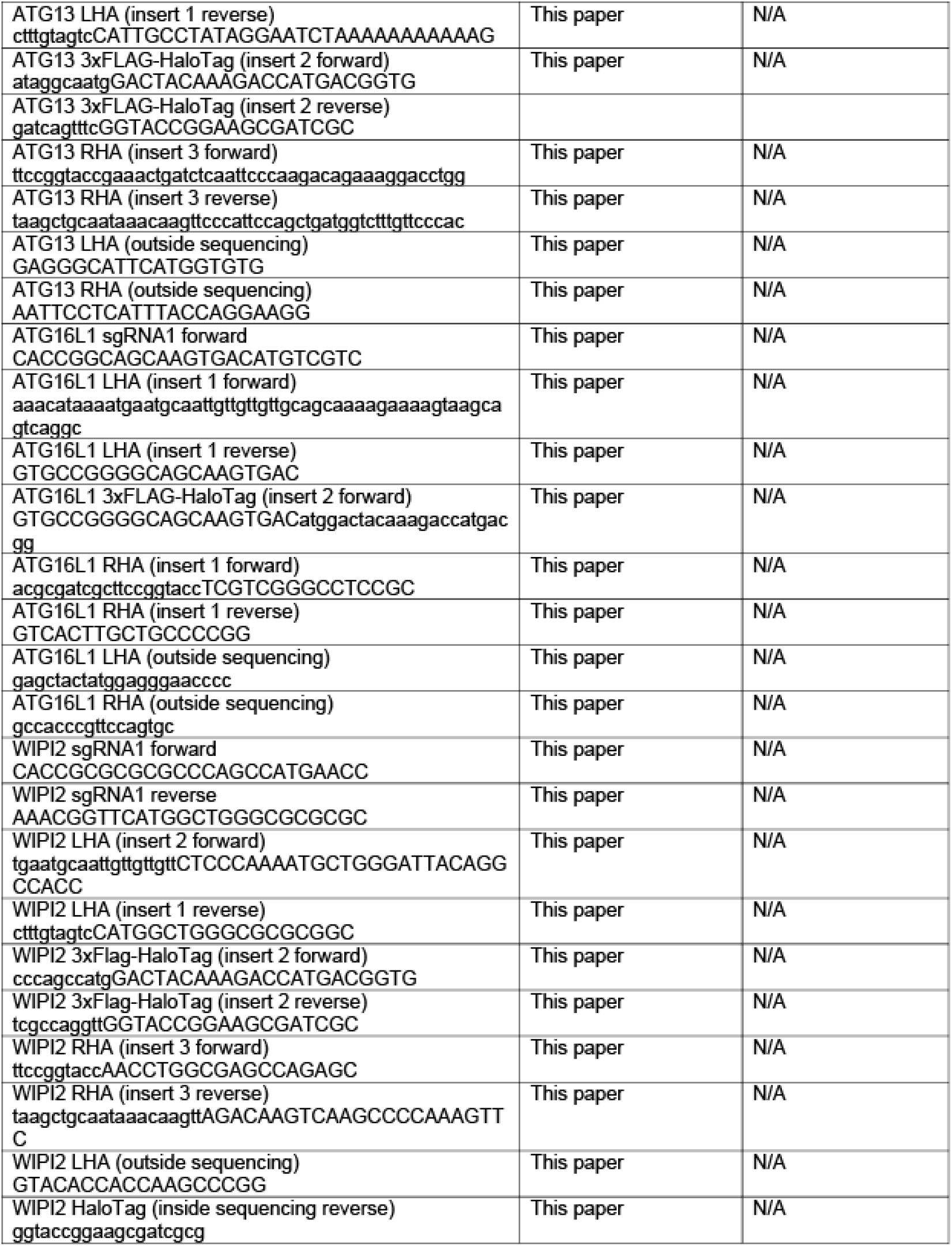
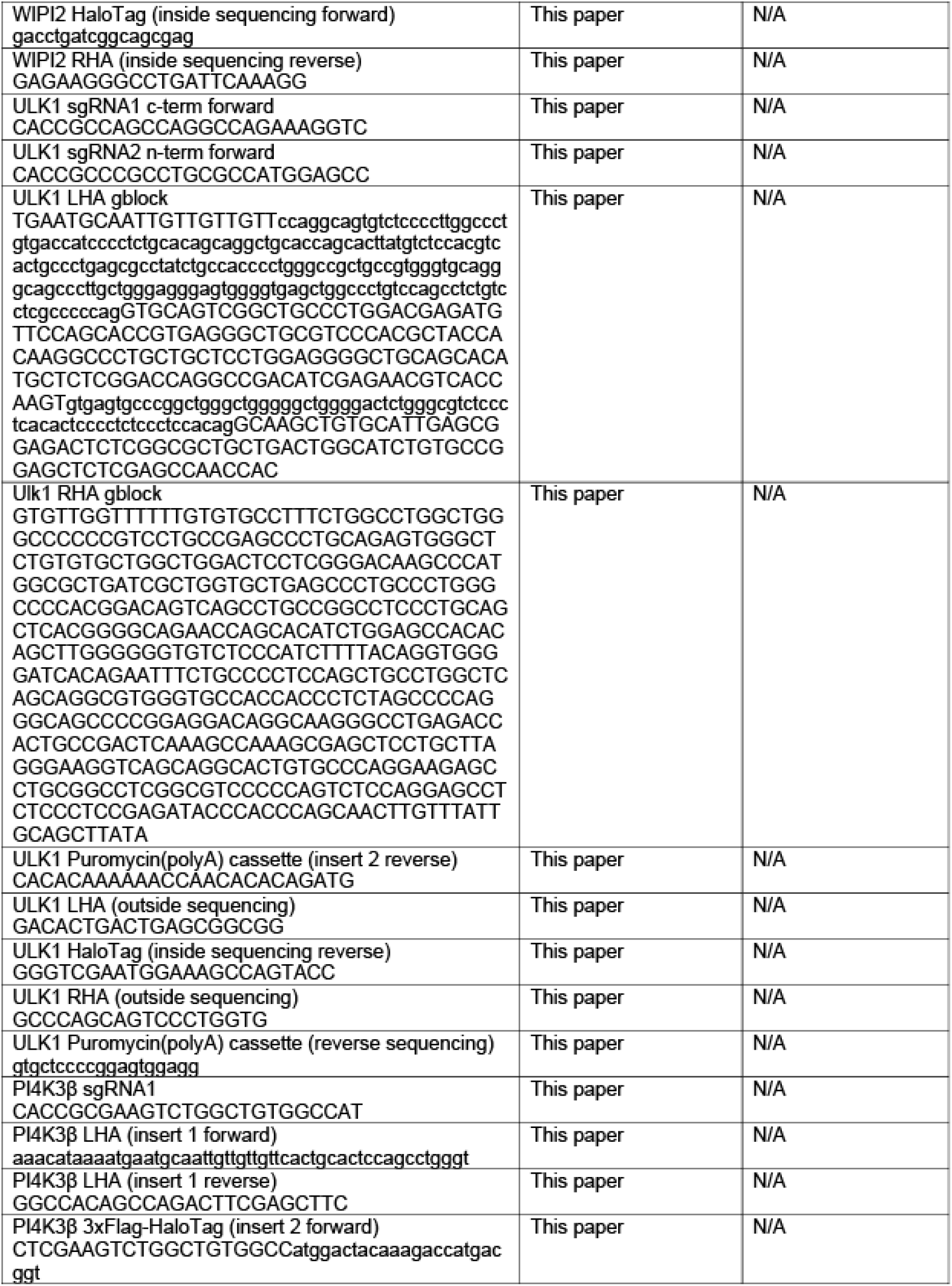
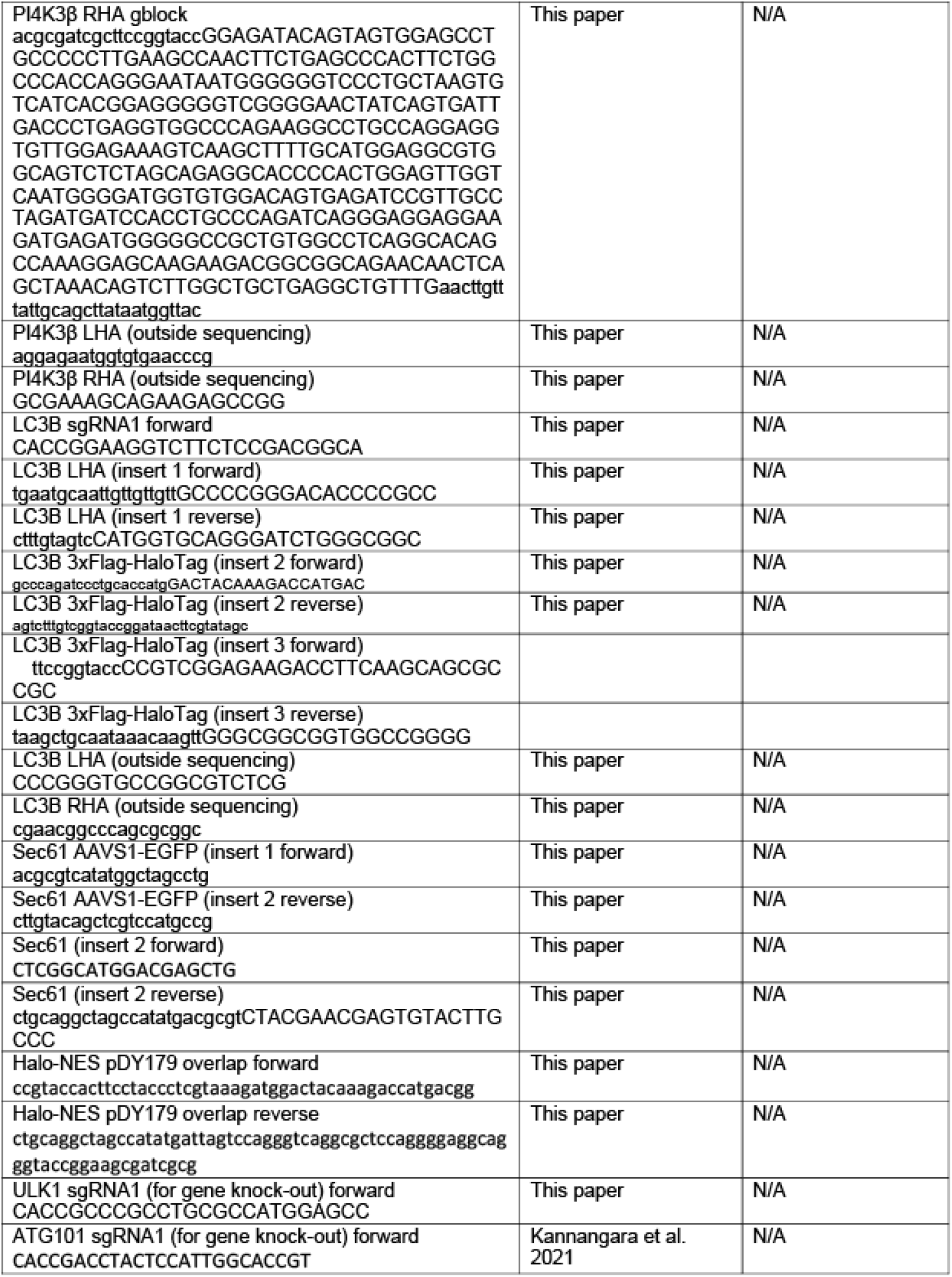
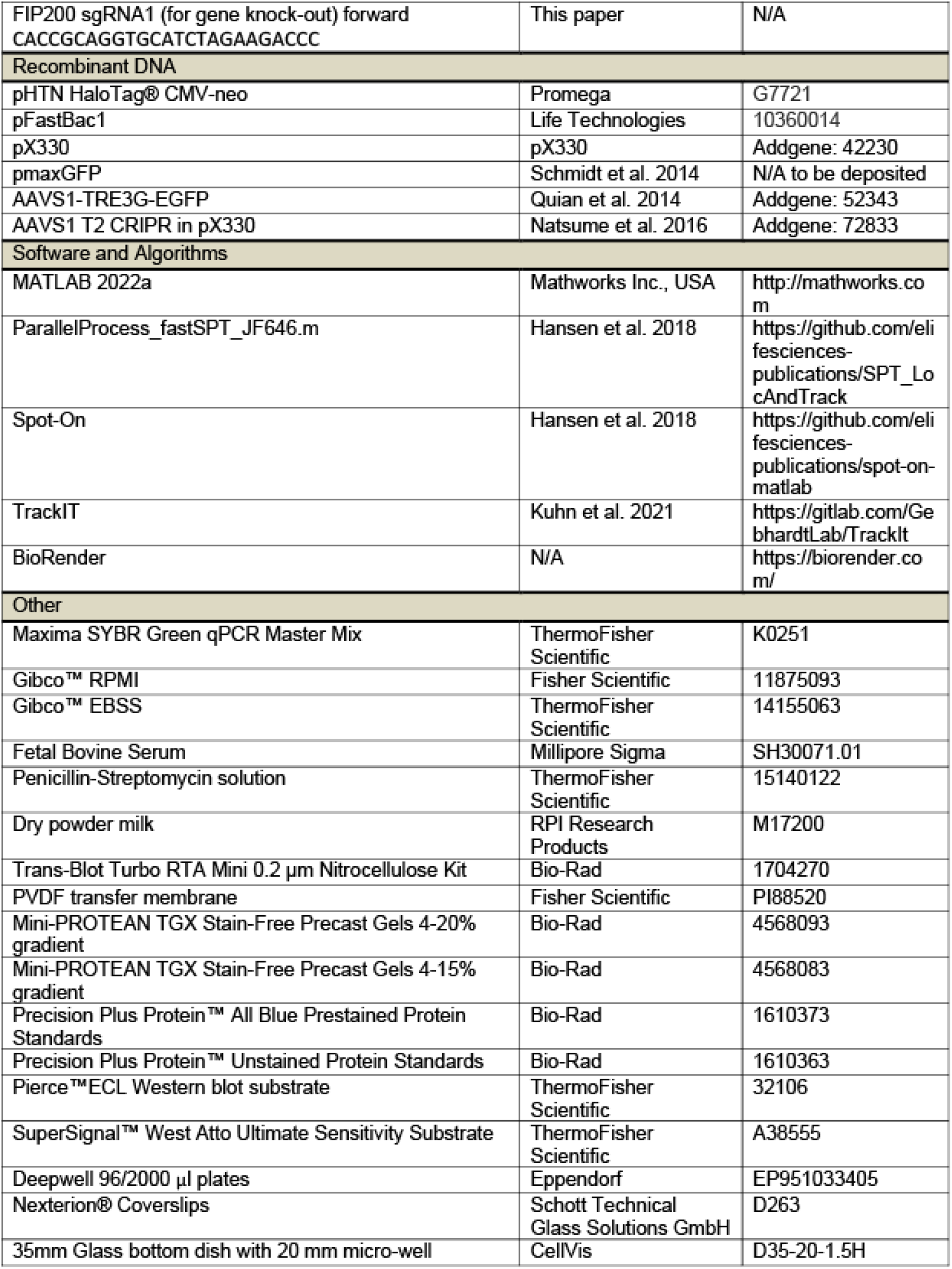

## References

Banerjee, C., Mehra, D., Song, D., Mancebo, A., Kim, D.-H., and Puchner, E.M. (2020). ULK1 forms distinct oligomeric states and nanoscopic morphologies during autophagy initiation. bioRxiv, 2020.2007.2003.187336. 10.1101/2020.07.03.187336.

Barth, S., Glick, D., and Macleod, K.F. (2010). Autophagy: assays and artifacts. J Pathol 221, 117–124. 10.1002/path.2694.

Berg, T.O., Fengsrud, M., Strømhaug, P.E., Berg, T., and Seglen, P.O. (1998). Isolation and characterization of rat liver amphisomes. Evidence for fusion of autophagosomes with both early and late endosomes. J Biol Chem 273, 21883–21892. 10.1074/jbc.273.34.21883.

Cattoglio, C., Pustova, I., Walther, N., Ho, J.J., Hantsche-Grininger, M., Inouye, C.J., Hossain, M.J., Dailey, G.M., Ellenberg, J., Darzacq, X., et al. (2019). Determining cellular CTCF and cohesin abundances to constrain 3D genome models. Elife 8. 10.7554/eLife.40164.

Chang, C., Jensen, L.E., and Hurley, J.H. (2021a). Autophagosome biogenesis comes out of the black box. Nat Cell Biol 23, 450–456. 10.1038/s41556-021-00669-y.

Chang, C., Shi, X., Jensen, L.E., Yokom, A.L., Fracchiolla, D., Martens, S., and Hurley, J.H. (2021b). Reconstitution of cargo-induced LC3 lipidation in mammalian selective autophagy. Sci Adv 7. 10.1126/sciadv.abg4922.

Chowdhury, S., Otomo, C., Leitner, A., Ohashi, K., Aebersold, R., Lander, G.C., and Otomo, T. (2018). Insights into autophagosome biogenesis from structural and biochemical analyses of the ATG2A-WIPI4 complex. Proc Natl Acad Sci U S A 115, E9792–E9801. 10.1073/pnas.1811874115.

Cong, L., Ran, F.A., Cox, D., Lin, S., Barretto, R., Habib, N., Hsu, P.D., Wu, X., Jiang, W., Marraffini, L.A., and Zhang, F. (2013). Multiplex genome engineering using CRISPR/Cas systems. Science 339, 819–823. 10.1126/science.1231143.

Dalle Pezze, P., Karanasios, E., Kandia, V., Manifava, M., Walker, S.A., Gambardella Le Novère, N., and Ktistakis, N.T. (2021). ATG13 dynamics in nonselective autophagy and mitophagy: insights from live imaging studies and mathematical modeling. Autophagy 17, 1131–1141. 10.1080/15548627.2020.1749401.

Dikic, I., and Elazar, Z. (2018). Mechanism and medical implications of mammalian autophagy. Nat Rev Mol Cell Biol 19, 349–364. 10.1038/s41580-018-0003-4.

Dooley, H.C., Razi, M., Polson, H.E., Girardin, S.E., Wilson, M.I., and Tooze, S.A. (2014). WIPI2 links LC3 conjugation with PI3P, autophagosome formation, and pathogen clearance by recruiting Atg12-5-16L1. Mol Cell 55, 238–252. 10.1016/j.molcel.2014.05.021.

Fracchiolla, D., Chang, C., Hurley, J.H., and Martens, S. (2020). A PI3K-WIPI2 positive feedback loop allosterically activates LC3 lipidation in autophagy. J Cell Biol 219. 10.1083/jcb.201912098.

Fujita, N., Hayashi-Nishino, M., Fukumoto, H., Omori, H., Yamamoto, A., Noda, T., and Yoshimori, T. (2008). An Atg4B mutant hampers the lipidation of LC3 paralogues and causes defects in autophagosome closure. Mol Biol Cell 19, 4651–4659. 10.1091/mbc.e08-03-0312.

Ganley, I.G., Lam, d.H., Wang, J., Ding, X., Chen, S., and Jiang, X. (2009). ULK1.ATG13.FIP200 complex mediates mTOR signaling and is essential for autophagy. J Biol Chem 284, 12297–12305. 10.1074/jbc.M900573200.

Ge, L., Zhang, M., Kenny, S.J., Liu, D., Maeda, M., Saito, K., Mathur, A., Xu, K., and Schekman, R. (2017). Remodeling of ER-exit sites initiates a membrane supply pathway for autophagosome biogenesis. EMBO Rep 18, 1586–1603. 10.15252/embr.201744559.

Gewirtz, D.A. (2014). The four faces of autophagy: implications for cancer therapy. Cancer Res 74, 647–651. 10.1158/0008-5472.CAN-13-2966.

Grimm, J.B., English, B.P., Chen, J., Slaughter, J.P., Zhang, Z., Revyakin, A., Patel, R., Macklin, J.J., Normanno, D., Singer, R.H., et al. (2015). A general method to improve fluorophores for live-cell and single-molecule microscopy. Nat Methods 12, 244–250, 243 p following 250. 10.1038/nmeth.3256.

Grimm, J.B., Muthusamy, A.K., Liang, Y., Brown, T.A., Lemon, W.C., Patel, R., Lu, R., Macklin, J.J., Keller, P.J., Ji, N., and Lavis, L.D. (2017). A general method to fine-tune fluorophores for live-cell and in vivo imaging. Nat Methods 14, 987–994. 10.1038/nmeth.4403.

Guardia, C.M., Tan, X.F., Lian, T., Rana, M.S., Zhou, W., Christenson, E.T., Lowry, A.J., Faraldo-Gómez, J.D., Bonifacino, J.S., Jiang, J., and Banerjee, A. (2020). Structure of Human ATG9A, the Only Transmembrane Protein of the Core Autophagy Machinery. Cell Rep 31, 107837. 10.1016/j.celrep.2020.107837.

Gómez-Sánchez, R., Rose, J., Guimarães, R., Mari, M., Papinski, D., Rieter, E., Geerts, W.J., Hardenberg, R., Kraft, C., Ungermann, C., and Reggiori, F. (2018). Atg9 establishes Atg2-dependent contact sites between the endoplasmic reticulum and phagophores. J Cell Biol 217, 2743–2763. 10.1083/jcb.201710116.

Hansen, A.S., Woringer, M., Grimm, J.B., Lavis, L.D., Tjian, R., and Darzacq, X. (2018). Robust model-based analysis of single-particle tracking experiments with Spot-On. Elife 7. 10.7554/eLife.33125.

Itakura, E., and Mizushima, N. (2010). Characterization of autophagosome formation site by a hierarchical analysis of mammalian Atg proteins. Autophagy 6, 764–776. 10.4161/auto.6.6.12709.

Judith, D., Jefferies, H.B.J., Boeing, S., Frith, D., Snijders, A.P., and Tooze, S.A. (2019). ATG9A shapes the forming autophagosome through Arfaptin 2 and phosphatidylinositol 4-kinase IIIβ. J Cell Biol 218, 1634–1652. 10.1083/jcb.201901115.

Kabeya, Y., Mizushima, N., Yamamoto, A., Oshitani-Okamoto, S., Ohsumi, Y., and Yoshimori, T. (2004). LC3, GABARAP and GATE16 localize to autophagosomal membrane depending on form-II formation. J Cell Sci 117, 2805–2812. 10.1242/jcs.01131.

Kannangara, A.R., Poole, D.M., McEwan, C.M., Youngs, J.C., Weerasekara, V.K., Thornock, A.M., Lazaro, M.T., Balasooriya, E.R., Oh, L.M., Soderblom, E.J., et al. (2021). BioID reveals an ATG9A interaction with ATG13-ATG101 in the degradation of p62/SQSTM1-ubiquitin clusters. EMBO Rep 22, e51136. 10.15252/embr.202051136.

Karanasios, E., Stapleton, E., Manifava, M., Kaizuka, T., Mizushima, N., Walker, S.A., and Ktistakis, N.T. (2013a). Dynamic association of the ULK1 complex with omegasomes during autophagy induction. J Cell Sci 126, 5224–5238. 10.1242/jcs.132415.

Karanasios, E., Stapleton, E., Walker, S.A., Manifava, M., and Ktistakis, N.T. (2013b). Live cell imaging of early autophagy events: omegasomes and beyond. J Vis Exp. 10.3791/50484.

Karanasios, E., Walker, S., Okkenhaug, H., Manifava, M., Hummel, E., Zimmermann, H., Ahmed, Q., Domart, M., Collinson, L., and Ktistakis, N. (2016). Autophagy initiation by ULK complex assembly on ER tubulovesicular regions marked by ATG9 vesicles. Nature Communications 7, ARTN 12420. 10.1038/ncomms12420.

Kirisako, T., Baba, M., Ishihara, N., Miyazawa, K., Ohsumi, M., Yoshimori, T., Noda, T., and Ohsumi, Y. (1999). Formation process of autophagosome is traced with Apg8/Aut7p in yeast. J Cell Biol 147, 435–446. 10.1083/jcb.147.2.435.

Kirkin, V. (2020). History of the Selective Autophagy Research: How Did It Begin and Where Does It Stand Today? J Mol Biol 432, 3–27. 10.1016/j.jmb.2019.05.010.

Kuhn, T., Hettich, J., Davtyan, R., and Gebhardt, J.C.M. (2021). Single molecule tracking and analysis framework including theory-predicted parameter settings. Sci Rep 11, 9465. 10.1038/s41598-021-88802-7.

Kuma, A., Matsui, M., and Mizushima, N. (2007). LC3, an autophagosome marker, can be incorporated into protein aggregates independent of autophagy: caution in the interpretation of LC3 localization. Autophagy 3, 323–328. 10.4161/auto.4012.

Kusama, Y., Sato, K., Kimura, N., Mitamura, J., Ohdaira, H., and Yoshida, K. (2009). Comprehensive analysis of expression pattern and promoter regulation of human autophagy-related genes. Apoptosis 14, 1165–1175. 10.1007/s10495-009-0390-2.

Lamb, C.A., Yoshimori, T., and Tooze, S.A. (2013). The autophagosome: origins unknown, biogenesis complex. Nat Rev Mol Cell Biol 14, 759–774. 10.1038/nrm3696.

Lin, M.G., and Hurley, J.H. (2016). Structure and function of the ULK1 complex in autophagy. Curr Opin Cell Biol 39, 61–68. 10.1016/j.ceb.2016.02.010.

Lu, J., Wu, M., and Yue, Z. (2020). Autophagy and Parkinson’s Disease. Adv Exp Med Biol 1207, 21–51. 10.1007/978-981-15-4272-52.

Lystad, A.H., Carlsson, S.R., de la Ballina, L.R., Kauffman, K.J., Nag, S., Yoshimori, T., Melia, T.J., and Simonsen, A. (2019). Distinct functions of ATG16L1 isoforms in membrane binding and LC3B lipidation in autophagy-related processes. Nat Cell Biol 21, 372–383. 10.1038/s41556-019-0274-9.

Martin, K.R., Celano, S.L., Solitro, A.R., Gunaydin, H., Scott, M., O’Hagan, R.C., Shumway, S.D., Fuller, P., and MacKeigan, J.P. (2018). A Potent and Selective ULK1 Inhibitor Suppresses Autophagy and Sensitizes Cancer Cells to Nutrient Stress. iScience 8, 74–84. 10.1016/j.isci.2018.09.012.

Matoba, K., Kotani, T., Tsutsumi, A., Tsuji, T., Mori, T., Noshiro, D., Sugita, Y., Nomura, N., Iwata, S., Ohsumi, Y., et al. (2020). Atg9 is a lipid scramblase that mediates autophagosomal membrane expansion. Nat Struct Mol Biol 27, 1185–1193. 10.1038/s41594-020-00518-w.

Matoba, K., and Noda, N.N. (2020). Secret of Atg9: lipid scramblase activity drives de novo autophagosome biogenesis. Cell Death Differ 27, 3386–3388. 10.1038/s41418-020-00663-1.

Mercer, C.A., Kaliappan, A., and Dennis, P.B. (2009). A novel, human Atg13 binding protein, Atg101, interacts with ULK1 and is essential for macroautophagy. Autophagy 5, 649–662. 10.4161/auto.5.5.8249.

Mercer, T.J., Gubas, A., and Tooze, S.A. (2018). A molecular perspective of mammalian autophagosome biogenesis. J Biol Chem 293, 5386–5395. 10.1074/jbc.R117.810366.

Mizushima, N. (2010). The role of the Atg1/ULK1 complex in autophagy regulation. Curr Opin Cell Biol 22, 132–139. 10.1016/j.ceb.2009.12.004.

Nakamura, S., and Yoshimori, T. (2017). New insights into autophagosome-lysosome fusion. J Cell Sci 130, 1209–1216. 10.1242/jcs.196352.

Nixon, R.A. (2007). Autophagy, amyloidogenesis and Alzheimer disease. J Cell Sci 120, 4081–4091. 10.1242/jcs.019265.

Noda, N.N. (2021). Atg2 and Atg9: Intermembrane and interleaflet lipid transporters driving autophagy. Biochim Biophys Acta Mol Cell Biol Lipids 1866, 158956. 10.1016/j.bbalip.2021.158956.

Ohashi, Y. (2021). Activation Mechanisms of the VPS34 Complexes. Cells 10. 10.3390/cells10113124.

Otomo, T., Chowdhury, S., and Lander, G.C. (2018). The rodshaped ATG2A-WIPI4 complex tethers membranes in vitro. Contact (Thousand Oaks) 1. 10.1177/2515256418819936.

Park, J.M., Jung, C.H., Seo, M., Otto, N.M., Grunwald, D., Kim, K.H., Moriarity, B., Kim, Y.M., Starker, C., Nho, R.S., et al. (2016). The ULK1 complex mediates MTORC1 signaling to the autophagy initiation machinery via binding and phosphorylating ATG14. Autophagy 12, 547–564. 10.1080/15548627.2016.1140293.

Ren, J., Liang, R., Wang, W., Zhang, D., Yu, L., and Feng, W. (2020). Multi-site-mediated entwining of the linear WIR-motif around WIPI β-propellers for autophagy. Nat Commun 11, 2702. 10.1038/s41467-020-16523-y.

Ren, X., Nguyen, T.N., Lam, W.K., Buffalo, C.Z., Lazarou, M., Yokom, A.L., and Hurley, J.H. (2022). Structural basis for ATG9A recruitment to the ULK1 complex in mitophagy initiation. bioRxiv, 2022.2007.2012.499634. 10.1101/2022.07.12.499634.

Russell, R.C., Tian, Y., Yuan, H., Park, H.W., Chang, Y.Y., Kim, J., Kim, H., Neufeld, T.P., Dillin, A., and Guan, K.L. (2013). ULK1 induces autophagy by phosphorylating Beclin-1 and activating VPS34 lipid kinase. Nat Cell Biol 15, 741–750. 10.1038/ncb2757.

Sawa-Makarska, J., Baumann, V., Coudevylle, N., von Bülow, S., Nogellova, V., Abert, C., Schuschnig, M., Graef, M., Hummer, G., and Martens, S. (2020). Reconstitution of autophagosome nucleation defines Atg9 vesicles as seeds for membrane formation. Science 369. 10.1126/science.aaz7714.

Schaaf, M.B., Keulers, T.G., Vooijs, M.A., and Rouschop, K.M. (2016). LC3/GABARAP family proteins: autophagy-(un)related functions. FASEB J 30, 3961–3978. 10.1096/fj.201600698R.

Schmidt, J.C., Zaug, A.J., and Cech, T.R. (2016). Live Cell Imaging Reveals the Dynamics of Telomerase Recruitment to Telomeres. Cell 166, 1188-1197.e1189. 10.1016/j.cell.2016.07.033.

Sergé, A., Bertaux, N., Rigneault, H., and Marguet, D. (2008). Dynamic multiple-target tracing to probe spatiotemporal cartography of cell membranes. Nat Methods 5, 687–694. 10.1038/nmeth.1233.

Shi, X., Yokom, A.L., Wang, C., Young, L.N., Youle, R.J., and Hurley, J.H. (2020). ULK complex organization in autophagy by a C-shaped FIP200 N-terminal domain dimer. J Cell Biol 219. 10.1083/jcb.201911047.

Shpilka, T., Weidberg, H., Pietrokovski, S., and Elazar, Z. (2011). Atg8: an autophagy-related ubiquitin-like protein family. Genome Biol 12, 226. 10.1186/gb-2011-12-7-226.

Stavoe, A.K., Gopal, P.P., Gubas, A., Tooze, S.A., and Holzbaur, E.L. (2019). Expression of WIPI2B counteracts age-related decline in autophagosome biogenesis in neurons. Elife 8. 10.7554/eLife.44219.

Szymańska, P., Martin, K.R., MacKeigan, J.P., Hlavacek, W.S., and Lipniacki, T. (2015). Computational analysis of an autophagy/translation switch based on mutual inhibition of MTORC1 and ULK1. PLoS One 10, e0116550. 10.1371/journal.pone.0116550.

Tang, Z., Takahashi, Y., He, H., Hattori, T., Chen, C., Liang, X., Chen, H., Young, M.M., and Wang, H.G. (2019). TOM40 Targets Atg2 to Mitochondria-Associated ER Membranes for Phagophore Expansion. Cell Rep 28, 1744-1757.e1745. 10.1016/j.celrep.2019.07.036.

Valverde, D.P., Yu, S., Boggavarapu, V., Kumar, N., Lees, J.A., Walz, T., Reinisch, K.M., and Melia, T.J. (2019). ATG2 transports lipids to promote autophagosome biogenesis. J Cell Biol 218, 1787–1798. 10.1083/jcb.201811139.

Waugh, M.G. (2019). The Great Escape: how phosphatidylinositol 4-kinases and PI4P promote vesicle exit from the Golgi (and drive cancer). Biochem J 476, 2321–2346. 10.1042/BCJ20180622.

White, E. (2015). The role for autophagy in cancer. Journal of Clinical Investigation 125, 42–46. 10.1172/JCI73941.

Yu, L., Chen, Y., and Tooze, S.A. (2018). Autophagy pathway: Cellular and molecular mechanisms. Autophagy 14, 207–215. 10.1080/15548627.2017.1378838.

